# Recurrent patterns of TOP1-mediated neuronal genomic damage shared by major neurodegenerative disorders

**DOI:** 10.1101/2025.03.03.641186

**Authors:** Zinan Zhou, Lovelace J. Luquette, Guanlan Dong, Junho Kim, Jayoung Ku, Kisong Kim, Nandini Ramesh, Mingyun Bae, Ann Caplin, Diane D. Shao, Bezawit Sahile, Kow Essuman, Eitan Goodman, Michael B. Miller, August Yue Huang, William J. Nathan, Andre Nussenzweig, Peter J. Park, Clotilde Lagier-Tourenne, Eunjung Alice Lee, Christopher A. Walsh

## Abstract

Amyotrophic lateral sclerosis (ALS), frontotemporal dementia (FTD), and Alzheimer’s disease (AD) represent two major categories of neurodegenerative disorders—TDP-43 and tau proteinopathies—for which the mechanisms driving neuronal death remain unclear. Single-cell whole-genome sequencing of 469 neurons from *C9ORF72* ALS, *C9ORF72* FTD, AD, and control brains revealed increased somatic single nucleotide variants (sSNVs) and insertions/deletions (sIndels) in all three diseases. Mutational signature analysis identified a disease-associated sSNV signature consistent with oxidative damage and an sIndel process affecting 22% of ALS, 76% of FTD, and 61% of AD neurons—but only 2% of control neurons—resembling signature ID4, previously linked to topoisomerase 1 (TOP1)-mediated mutagenesis. Rapid Approach to DNA Adduct Recovery (RADAR) assays confirmed increased TOP1-DNA covalent complexes and duplex sequencing confirmed the increased sIndels and identified single-strand events as likely precursor lesions. TOP1-associated sIndel mutagenesis and genome instability thus represents a mechanism shared by both TDP-43 and tau neurodegeneration.

## INTRODUCTION

Amyotrophic lateral sclerosis (ALS), frontotemporal dementia (FTD) and Alzheimer’s disease (AD)^1,2^ are common neurodegenerative diseases with diverse pathologies and genetic underpinnings. ALS and FTD are closely related, with clinical, pathological, and genetic overlap, marked by the depletion of nuclear TDP-43 and accumulation of cytoplasmic TDP-43 inclusions in neurons. Mutations in many genes, such as *C9ORF72*, *TARDBP (TDP-43)*, *FUS, TBK1, VCP,* and *SQSTM1* contribute to both diseases^3^, with *C9ORF72* repeat expansion being the most common genetic cause of both familial and sporadic forms of ALS and FTD^4,5^. In contrast, AD pathology is characterized by accumulation of amyloid-β protein and phosphorylated tau, prominent activation of microglia, and shows distinct underlying genetic risks^6^.

Genomic instability, which leads to the accumulation of DNA damage and mutations, has been implicated in all of these conditions, but its exact role remains elusive. In ALS, FTD and AD, increased DNA damage due to high levels of oxidative stress, R-loops and DNA strand breaks has been reported^7–16^. Several ALS/FTD-related genes are involved in DNA damage responses, such as *SOD1*, *FUS*, *TDP-43*, *C9ORF72*, *NEK1*, *SQSTM1*, *SETX,* and *VCP*^9,16–24^. *C9ORF72* repeat expansions have recently been shown to induce chromosomal fragility and DNA damage^25^. Neurons are particularly vulnerable to DNA damage due to their high transcriptional activity, large energy demand and associated oxidative stress. Recent studies have shown that somatic single nucleotide variants (sSNVs) accumulate linearly with age in neurons, with further increases observed in neurodegenerative conditions, including AD, Cockayne syndrome, and Xeroderma pigmentosum^13,14,26^.

In the present study, we compared somatic mutations in neurons from major neurodegenerative diseases associated with TDP-43 proteinopathy (ALS and FTD) and tau proteinopathy (AD) to compare potential mechanisms of mutagenesis between these two major categories of neurodegenerative diseases. We profiled sSNVs and somatic insertions and deletions (sIndels) in neurons with and without depletion of nuclear TDP-43 from the motor or premotor cortex of six *C9ORF72* ALS and the prefrontal cortex of six *C9ORF72* FTD brains using single-cell whole-genome sequencing (scWGS). We then compared the results to those from AD neurons^14^, in which sIndels were not analyzed, and neurons from neurotypical controls. Our analysis revealed significantly elevated levels of sSNVs and sIndels in neurons from *C9ORF72* ALS, *C9ORF72* FTD and AD brains, but in surprisingly similar patterns despite their diverse genetic basis and pathologies. Importantly, our findings indicate that neurons in all three disease conditions often exhibit an excessive number of sIndels—sometimes over 1000, equivalent to hundreds of years of age-related sIndel accumulation—and share a mutational pattern characterized by two-basepair (2-bp) deletions. Computational and experimental analyses of these sIndels associate the mutagenic process to topoisomerase 1 (TOP1)-mediated mutagenesis, and show that this mutagenic process is absent in cerebellar neurons from the same diseased brains. Our results suggest that increased DNA damage and accelerated somatic mutation contribute to the pathogenesis of degenerative disorders with diverse pathological features.

## RESULTS

### Isolation of neuronal nuclei with TDP-43 pathology from *C9ORF72* ALS and FTD brains achieves high purity

To isolate neuronal nuclei with either normal or depleted nuclear TDP-43 (TDP-43+ and TDP-43-) from postmortem *C9ORF72* ALS and FTD brains, we performed fluorescence-activated nuclear sorting (FANS) with co-staining of NeuN and TDP-43 antibodies^27^ (Figure 1A). TDP-43- neurons were only found in *C9ORF72* ALS and FTD brains and were absent in age-matched controls (Figure 1B). Single-nucleus RNA sequencing (snRNA-seq) data from isolated nuclei revealed that 96.4% of TDP-43+/NeuN+ and 95.4% of TDP-43-/NeuN+ nuclei were indeed neuronal (Figures 1C and S1A). Furthermore, we examined cryptic exons caused by loss of TDP-43 function during RNA splicing in the snRNA-seq data^28–30^. Transcripts containing known cryptic exons in the *RAP1GAP*, *STMN2*, *ATP8A2,* and *KALRN* genes were detected in TDP-43- neurons but were absent or very rare in TDP-43+ neurons (Figures 1D, 1E, S1B and S1C), confirming the presence of TDP-43 disease pathology in isolated TDP-43- neurons.

**Figure 1.**
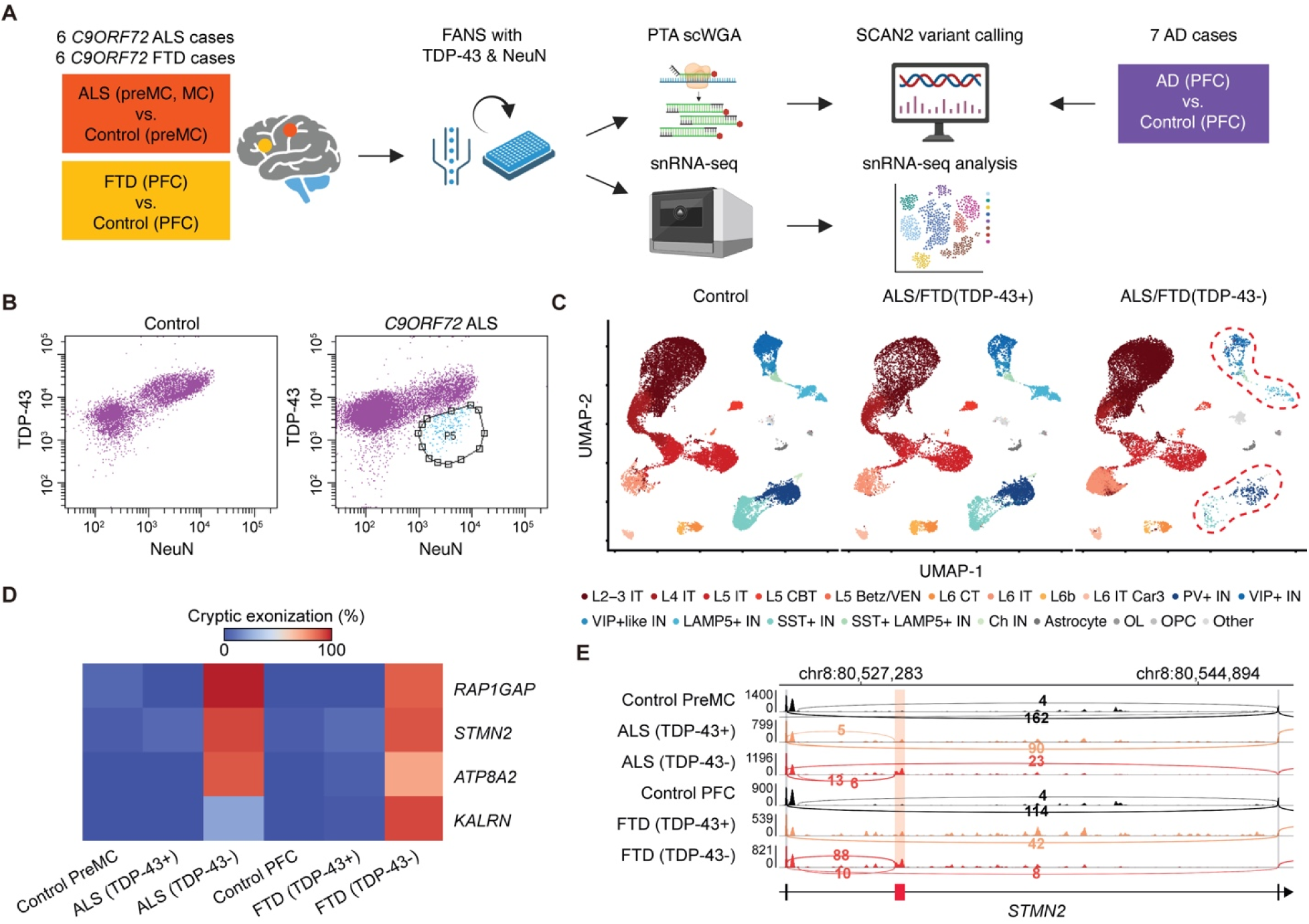
Experimental strategy and characterization of isolated neuronal nuclei from *C9ORF72* ALS and FTD brains. (A) Overview of the experimental design. Neuronal nuclei with and without nuclear TDP-43 depletion were isolated from 6 *C9ORF72* ALS and 6 *C9ORF72* FTD brains, then subjected to genome sequencing and analysis. These were compared with neuronal nuclei from neurotypical brains, including newly generated data and previously published datasets. Previously generated scWGS data from AD brains were included as well. (B) Representative FACS plots showing neuronal nuclei from a *C9ORF72* ALS brain and a neurotypical control brain co-stained with NeuN and TDP-43 antibodies. The P5 gate (right, circle) indicates neuronal nuclei with depletion of nuclear TDP-43. (C) UMAP clustering of snRNA-seq data from neuronal nuclei, colored by major cell types. TDP-43- neuronal nuclei come exclusively from *C9ORF72* ALS and FTD brains, while TDP-43+ neuronal nuclei come from both diseased and neurotypical brains; control indicates neurotypical brains. IT: intratelencephalic neurons. CBT: corticobulbar tract neurons. VEN: Von Economo neurons. CT: corticothalamic neurons. PV: parvalbumin. IN: inhibitory neurons. Ch IN: cholinergic inhibitory neurons. OL: oligodendrocytes. OPC: oligodendrocyte precursor cells. Red dashed circles emphasize reduced inhibitory neurons under the TDP-43- condition (D) Heatmap of cryptic exonization showing the proportion of transcripts with cryptic exons in neurons across different conditions. (E) IGV Genome browser tracks of *STMN2* showing snRNA-seq read coverage and splice junctions. TDP-43- neurons (red) contain a marked increase in cryptic exon inclusion (highlighted region), which is absent or low in TDP-43+ (orange) and control (black) neurons. Numbers next to splice junctions are read counts supporting the junction. PFC: prefrontal cortex. preMC: premotor cortex. See also Figure S1

Both TDP-43+ and TDP-43- neurons encompassed all major clusters of excitatory and inhibitory neurons (Figure 1C), though inhibitory neurons were notably depleted in the TDP-43-population (Figures 1C and S1D), confirmed by the reduced expression of inhibitory neuron markers in TDP-43- neurons compared to TDP-43+ neurons in previously generated bulk RNA-seq data from *C9ORF72* FTD brains (Figure S1E)^27^. These results are consistent with recent studies demonstrating that inhibitory neurons rarely exhibit cytoplasmic TDP-43 inclusions^31^. However, it remains unclear whether the reduced TDP-43 pathology in inhibitory neurons reflects resistance or susceptibility to TDP-43 pathology given the observed loss of inhibitory neurons in ALS^32–34^.

### Somatic mutation burdens are increased in diseased neurons

We performed primary template-directed amplification (PTA)-based single-cell whole-genome amplification (scWGA) and sequencing (scWGS) on neurons isolated from the motor cortex or premotor cortex of six *C9ORF72* ALS brains and the prefrontal cortex of six *C9ORF72* FTD brains (three TDP-43+ and three TDP-43- neurons per brain; Tables S1 and S2), and identified sSNVs and sIndels using Single Cell ANalysis 2 (SCAN2)^26^, our recently developed single-cell genotyper (Figure 1A). Somatic mutations in these diseased neurons were compared with those in 24 newly sequenced premotor cortex neurons and 56 previously sequenced prefrontal cortex neurons from neurotypical controls across various ages^26,35^, as well as 29 previously sequenced prefrontal cortex neurons from AD brains (Figure 1A and Tables S1, S2 and S3)^14^. Cells that did not pass quality control (QC) criteria, such as amplification uniformity^36^, were excluded from downstream analyses (Figures S2, S3 and Table S2, see STAR Methods). SCAN2 recovered approximately 35% of sSNVs and 24% of sIndels, producing a somatic mutation catalog with >67,000 sSNVs and >12,000 sIndels with an aggregate estimated error rate of <10% (Figure S2 and Table S4).

Many neurons from *C9ORF72* ALS, *C9ORF72* FTD and AD brains exhibited elevated burdens of both sSNVs and sIndels, with sIndels showing greater increases (Figure 2A and Table S5). Substantial variability in sSNV and sIndel burdens were observed between neurons from brains with the same disease, and in some cases even within the same brain (Data S1). Since age-matched neurons from the prefrontal and premotor cortex of neurotypical controls aged >50 years showed similar burdens of sSNVs and sIndels (Figure 2B) neurons from these two sites were combined in all subsequent analyses. After adjusting for age, burdens of both sSNVs and sIndels in neurons from FTD and AD were significantly increased compared to those from neurotypical controls (Figure 2C). In ALS neurons, sIndel burdens were significantly increased, but not sSNV burdens (Figure 2C). The relatively modest increase in somatic mutations in *C9ORF72* ALS neurons compared to *C9ORF72* FTD and AD neurons may reflect the selective involvement of motor neurons in ALS, and their rapid and almost complete disappearance by the time of death, whereas a broader range of neuronal types are involved in FTD and AD^37^. Together, our results show that accumulation of sSNVs and sIndels is accelerated in *C9ORF72* ALS, *C9ORF72* FTD and AD neurons. Unexpectedly, TDP-43- neurons from *C9ORF72* ALS and FTD brains did not exhibit higher burdens of sSNVs and sIndels compared to TDP-43+ neurons (Figures 2D), despite TDP-43’s involvement in DNA damage repair of double-strand breaks^9^. Merging neurons from *C9ORF72* ALS and FTD cases to increase statistical power suggested a marginal increase in sIndels in TDP-43+ relative to TDP-43- neurons (Figure 2E), but this effect was small and may reflect differences in neuronal survival or sampling. Nonetheless, our data suggest that *C9ORF72* repeat expansion and depletion of nuclear TDP-43 are not synergistic in causing DNA damage.

**Figure 2.**
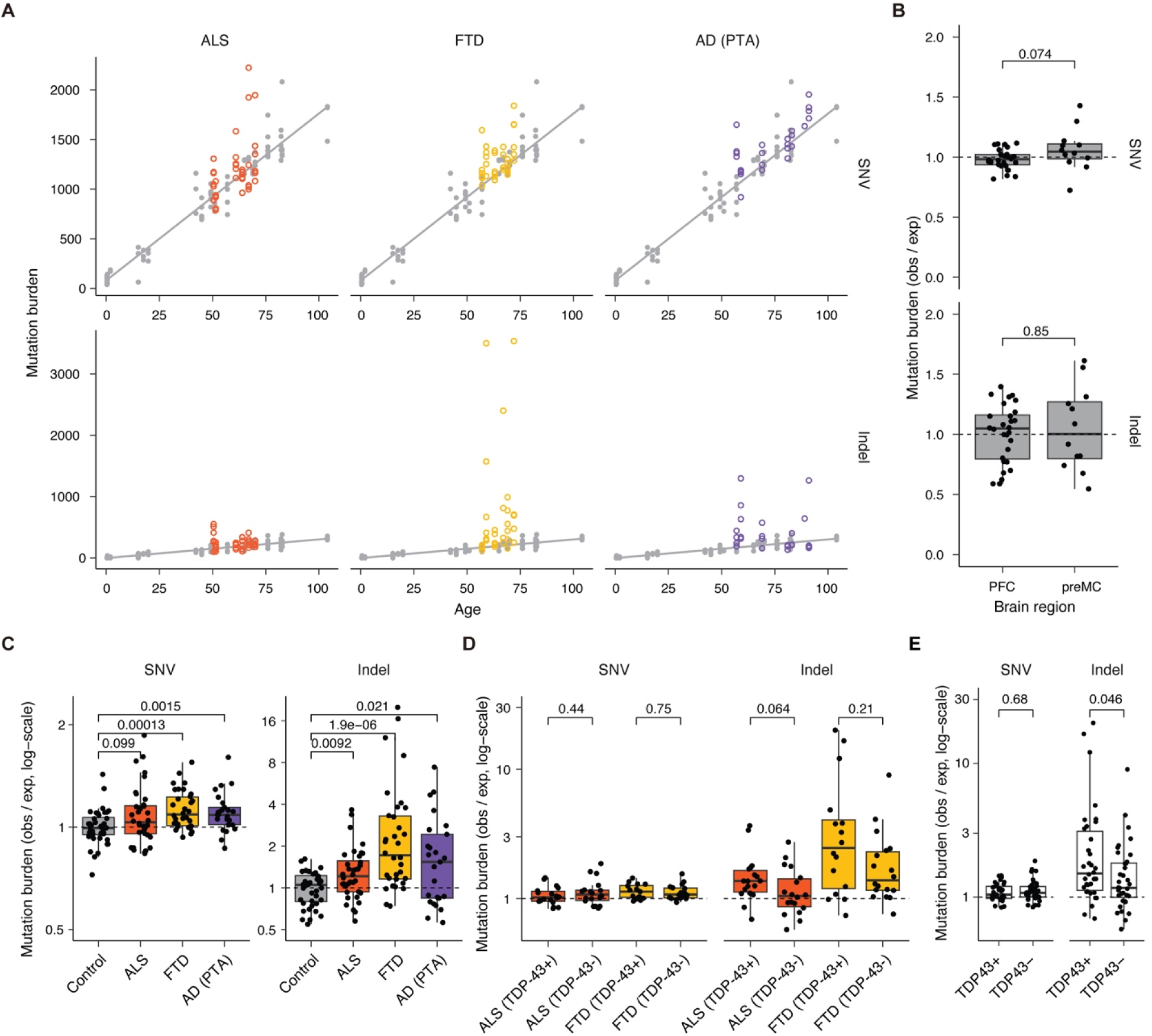
Increased somatic mutation burdens in neurons from *C9ORF72* ALS, *C9ORF72* FTD and AD brains. (A) Extrapolated genome-wide sSNV and sIndel burdens for neurons from *C9ORF72* ALS, *C9ORF72* FTD and AD brains compared to those for neurons from neurotypical brains as a function of age. Identical control neuron points and regression (see STAR Methods) are shown in gray in each panel for comparison. (B) Burdens of sSNVs and sIndels in PFC (prefrontal cortex) and preMC (premotor cortex) neurons from neurotypical brains aged >50 years. Each point represents the burden of one neuron, compared against the expected burden for its age (see STAR Methods). (C) Mutation burdens corrected for age (see STAR Methods); preMC and PFC neurons are combined to form the control set. (D) TDP-43+ and TDP-43- neurons from *C9ORF72* ALS and FTD brains show similar levels of sSNVs and sIndels. All *P*-values in this figure represent Wilcoxon rank-sum tests. (E) Neurons from ALS and FTD cases were combined into a single group of TDP43+ and TDP43- neurons. *P*-values are two-sided Wilcoxon rank-sum tests and none are corrected for multiple hypothesis tests as they already are not significant. See also Figures S2 and -S3

### A recurrent pattern of 2-bp deletions across diseased neurons

Among several patterns of mutations observed in our mutation catalog, we uncovered a disease-specific signature characterized by 2-bp deletions that was greatly elevated in neurons from all three disease conditions compared to controls (Figures 3A and 3B and Table S5). To investigate potential mutagenic mechanisms, we performed *de novo* mutational signature extraction, a technique that can identify underlying processes based on the types and frequencies of mutations^38^. *De novo* mutational signature extraction was performed on neuronal mutations from the three neurodegenerative conditions, along with (1) 81 neurons from AD, as well as 190 neurons and 40 glial cells from neurotypical controls previously amplified using multiple displacement amplification (MDA), an earlier whole-genome amplification method^39^, and (2) an additional 66 oligodendrocytes from neurotypical controls amplified using PTA^35^ This analysis resulted in six *de novo* SNV signatures (SBS-A to SBS-F, Figure S4A) of which only SBS-B showed an increase in PTA neurons from all three neurodegenerative conditions compared to control PTA neurons, although the increase in ALS neurons did not reach statistical significance (Figures S4B and S4C).

**Figure 3.**
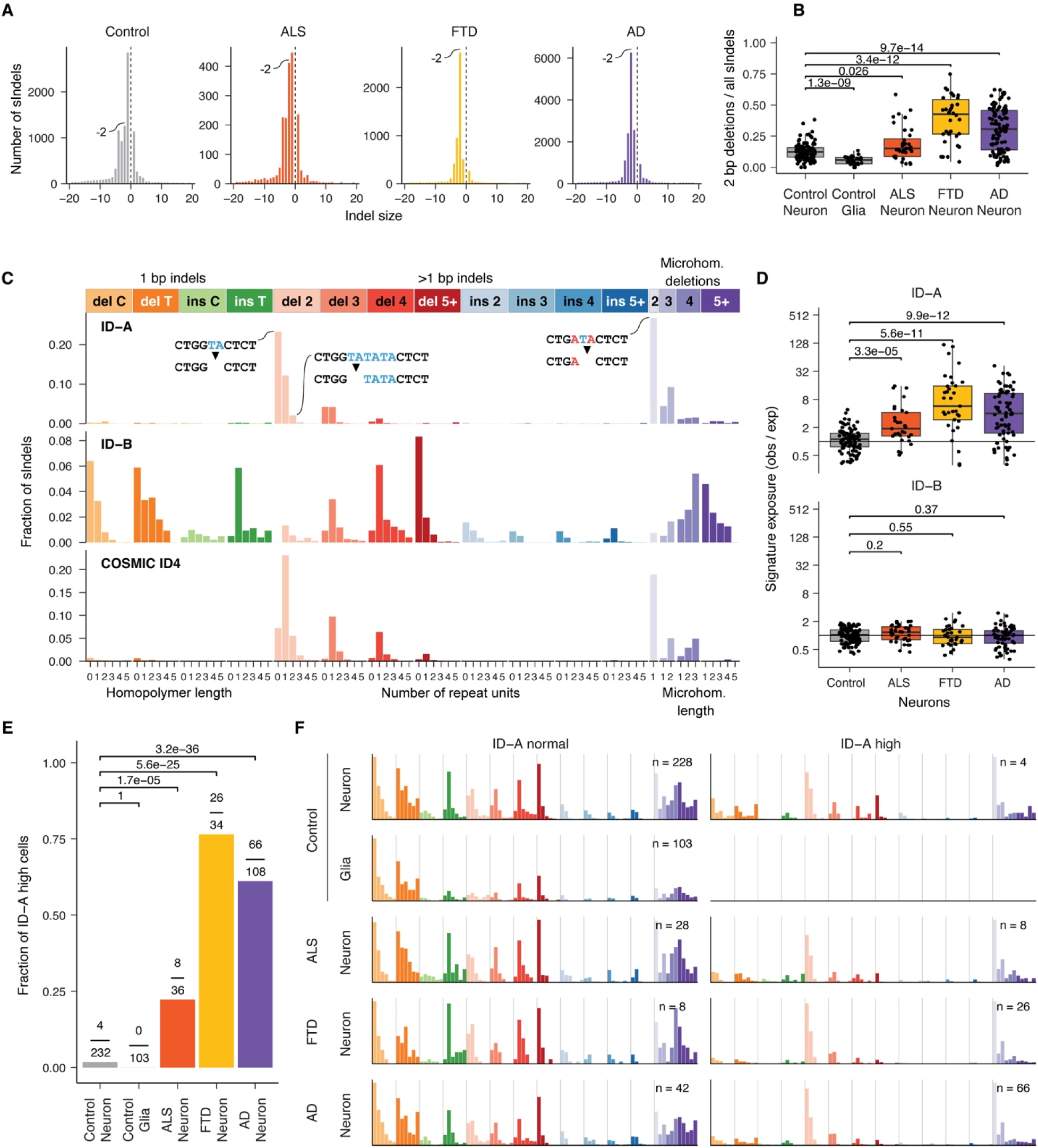
Neurons from *C9ORF72* ALS, *C9ORF72* FTD and AD brains share an Indel mutational signature. (A) Distribution of sIndel lengths across conditions. Diseased neurons have higher levels of 2-bp deletions compared to controls. (B) Fraction of 2-bp deletions in neurons and glia across conditions. *P*-values represent Wilcoxon rank-sum tests. Glia include pure oligodendrocytes and mixed glia. (C) Two *de novo* sIndel signatures identified from joint analysis of diseased and neurotypical control neurons amplified by PTA, as well as previously published neurons and glia amplified using an older amplification technology (MDA). COSMIC ID4 is a signature previously identified in cancer and normal aging neurons. Three examples of deletions that would be classified into the corresponding bars are provided to illustrate the disease-associated 2-bp deletions in ID-A. (D) Burdens of ID-A and ID-B sIndels; *P*-values represent Wilcoxon rank-sum tests. (E) Fraction of cells with high levels of ID-A sIndels in neurons and glia across conditions, determined by a classifier (see STAR Methods); *P*-values represent Fisher tests of independence between disease status and ID-A-high classifications. Two neurons and three oligodendrocytes with 0 sIndels were excluded from classification. (F) sIndel spectra of neurons and glia stratified by ID-A-high and -normal classifications. See also Figures S5 and S6

To explore the mutagenic processes associated with the *de novo* signatures, we compared them to existing mutational signatures in the Catalogue of Somatic Mutations in Cancer (COSMIC) database. SBS-B closely resembles the COSMIC SBS30 signature (cosine similarity 0.90), which is associated with inactivating mutations in *NTHL1,* an enzyme involved in base excision repair of oxidative DNA damage^40^. The elevation of SBS-B sSNVs in neurons from neurodegenerative conditions may therefore indicate increased levels of oxidative DNA damage, consistent with previous studies revealing increased oxidative stress in these diseases^7,10,12^. SBS-A and SBS-D resemble COSMIC SBS5 and SBS1, respectively, both of which are “clock-like” signatures. While SBS1 is thought to be cell-division dependent and thus limited in postmitotic neurons, SBS5 accumulates universally during aging. The mechanisms underlying SBS-C and SBS-E (which showed no increase in diseased neurons) and SBS-F (which was increased in ALS and AD neurons) are unclear.

*De novo* mutational signature extraction of sIndels identified three distinct signatures, ID-A, ID-B, and ID-C (Figure 3C and S5A). ID-A burdens were significantly increased in PTA neurons from the three neurodegenerative conditions compared to control PTA neurons, while burdens of ID-B showed no difference across PTA neurons from all conditions (Figures 3D and S5B-C). ID-C was elevated only in MDA-amplified AD neurons, but not in PTA-amplified AD neurons (*p* = 3.9 × 10^-4^, Wilcoxon rank sum test), suggesting that this signature may represent overfitting to MDA-specific artifacts (Figure S5C). ID-A is dominated by 2-bp deletions (Data S2) and most closely resembles the COSMIC signature ID4 (cosine similarity 0.80), which has recently been linked to deficiency in RNase H2, an enzyme responsible for removing ribonucleotides erroneously incorporated into the genome. In the absence of functional RNase H2, ribonucleotides can instead be removed through TOP1-mediated ribonucleotide excision repair (RER)^41^. Although ID4 is a feature of normal aging in neurotypical neurons—even more so than the canonical “clock-like” Indel signatures ID5 and ID8 identified in cancers^26,35,42^—the excessive ID-A burden observed in all three neurodegenerative conditions suggests a dramatic acceleration of this age-associated process. Indeed, virtually all of the excess sIndel burdens in diseased neurons can be attributed to ID-A.

We constructed a classifier (see STAR Methods) to determine which cells exhibited ID-A burdens in line with typical neuronal aging (ID-A normal) or an excessive ID-A burden (ID-A high). Upon stratifying our cells by ID-A status (Figure 3E), we found significantly higher fractions of ID-A high neurons in every neurodegenerative condition (ALS: OR = 15.1, *p* = 1.68 × 10^-5^; FTD: OR = 158.3, *p* = 5.58 × 10^-25^; AD: OR = 79.5, *p* = 3.22 × 10^-36^; all comparisons represent one-sided Fisher’s tests vs. control neurons) while only 4/232 control neurons and none of the 103 control glia were classified as ID-A high. Furthermore, 14 of 19 diseased brains contained at least one ID-A-high PTA-amplified neuron, and the mutational spectra of ID-A-high and -normal neurons were remarkably similar across conditions (Figure 3F), supporting the notion that ID-A pathology is likely a universal feature of these diverse neurodegenerative conditions rather than an isolated occurrence. Consistent with the overall burdens of sSNVs and sIndels, the levels of these *de novo* mutational signatures did not differ between TDP-43+ and TDP-43- neurons (Figure S6), further indicating that nuclear TDP-43 depletion does not enhance mutagenesis in individuals with *C9ORF72* expansions.

### ID-A deletions exhibit features of TOP1-mediated mutagenesis

To further investigate the association between ID-A sIndels in diseased neurons and TOP1-mediated mutagenesis, we examined features of the 2-bp deletions. Mammalian TOP1 exhibits a preference for thymine bases and cleaves at the 3’ phosphodiester bond following the thymine nucleotide^43^, primarily causing deletions at a TNT motif, with the most common deletions being CT dinucleotides (Figure 4A)^41^. Consistent with this observation, the 2-bp deletions in neurons from neurodegenerative conditions were enriched in CT deletions and deletions with microhomology or at least 2 repeat units occurred most frequently within TNT motifs (Figures 4B and 4C).

**Figure 4.**
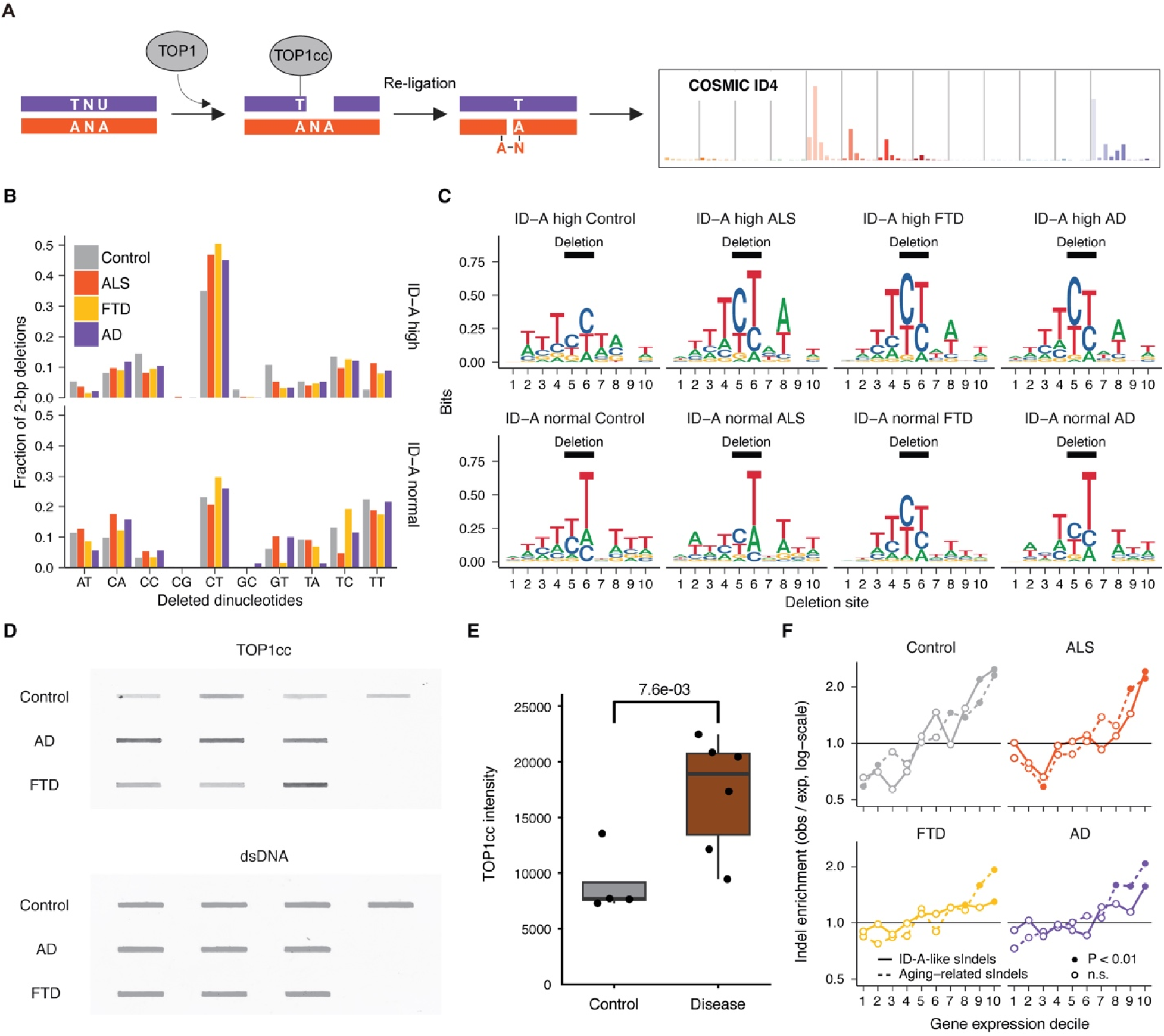
Disease-associated ID-A-like deletions reflect characteristics of TOP1-mediated ribonucleotide excision repair. (A) Model of TOP1-mediated ribonucleotide excision repair. (B) Fraction of 2-bp deletions by the deleted dinucleotide stratified by ID-A-high and -normal classifications. (C) DNA motifs showing the ten basepair context around 2-bp deletions with at least 2 repeat units and stratified by ID-A-high and -normal classifications; positions 5 and 6 are the deleted nucleotides. (D) RADAR assay measuring TOP1 cleavage complex (TOP1cc) levels in NeuN+ neuronal nuclei from *C9ORF72* FTD, AD and control brains. Slot blots were probed with antibodies against TOP1cc and dsDNA. Each band represents one individual. (E) Quantification of TOP1cc levels in arbitrary units in (D) using ImageJ. Each dot represents one individual. The *P*-value represents an unpaired one-sided Welch’s *t*-test. (F) Enrichment of sIndels in relation to local gene expression levels from previously published snRNA-seq data of excitatory neurons. Enrichment tests divide the genome into 10 roughly equally-sized, non-contiguous regions ranked by gene expression. I.e., decile 10 represents the subset of the genome containing the top 10% of expressed genes. Solid lines: enrichment analysis of ID-A-like sIndels (see Figure 3C), dotted lines: all other sIndels. *P*-values are enrichment tests (see STAR Methods) indicating that the observed/expected ratio significantly differs from 1. See also Figures S7 and S8

Neurons from *C9ORF72* FTD and AD cases further showed increased levels of the TOP1 cleavage complex (TOP1cc), a protein complex formed through a covalent TOP1-DNA crosslink during TOP1-mediated RER (Figure 4A).We performed the Rapid Approach to DNA Adduct Recovery (RADAR) assay to measure TOP1cc levels directly in neuronal nuclei isolated from the prefrontal cortex of *C9ORF72* FTD, AD, and age-matched control brains^44^. TOP1cc signals were significantly increased in diseased neurons compared to controls (Figures 4D and 4E, see STAR Methods). In parallel, we analyzed neuronal nuclei that were positive for both TOP1cc and NeuN antibody staining from the prefrontal cortex of *C9ORF72* FTD, AD, and control brains using flow cytometry and found higher fractions of neurons with increased TOP1cc immunoreactivity in disease brains (2.8-11.6%) compared to controls (0.2-0.5%) (Figure S7A). These results provide direct evidence of increased formation of covalent TOP1 links to DNA in *C9ORF72* FTD and AD neurons.

TOP1-mediated ID4 Indels can arise during both replication and transcription in mitotic cells^45^, and are enriched at transcribed regions in cells with RNase H2 deficiency^41^. In normal postmitotic neurons, ID4 Indels are also strongly associated with transcription^35^. To explore the association between sIndels and transcription, we first split all sIndels into two categories: (1) all 2-bp deletions and the 2-bp microhomology deletions that characterize ID-A (Figure 3C) and (2) all other sIndels. We then compared the burdens of each sIndel category to local gene expression levels measured in excitatory neurons from previously published snRNA-seq data^35^. In neurotypical control neurons, both sIndel categories were strongly associated with expression levels (Figure 4F). In the neurodegenerative conditions (which contain higher amounts of ID-A-like sIndels), both categories remained positively associated with expression, though the association was weaker for ID-A-like sIndels. This association with transcriptional activity was confirmed by comparing each sIndel category against several other signals that correlate with transcriptional activity: local levels of open chromatin (measured by single-nucleus ATAC-seq^35^), active and inactive histone marks^46^, chromatin states^47^, and promoters and enhancers active in neurons^48^ (Figures S7B and S7C). Taken together, these characteristics of TOP1-related Indel mutagenesis suggest that the ID-A-like sIndels observed in our diseased neurons are likely TOP1-mediated in these three neurodegenerative conditions.

### Single-strand breaks and deletions are prevalent in neurodegenerative conditions

The removal of incorporated ribonucleotides during TOP1-mediated RER creates single-strand breaks (Figure 4A), and analysis of genomic DNA (gDNA) showed directly that cortex tissues from diseased cases with large numbers of sIndels exhibited widespread DNA fragmentation. To assess these single-strand breaks, gDNA extracted from disease-relevant regions of *C9ORF72* ALS, *C9ORF72* FTD, AD, and neurotypical control brains was treated with S1 nuclease and analyzed using agarose gel electrophoresis (Figure 5A, see STAR Methods). S1 nuclease converts single-strand breaks into double-strand breaks, which appear as smears on the gel. We then quantified smear intensity using TapeStation and found significantly elevated levels of fragmented single-strand gDNA in diseased brains (Figure 5B), with no correlation to donor age (Figure 5C). Furthermore, the levels of fragmented single-strand gDNA in diseased brains were significantly correlated with the burdens of ID-A sIndels, accounting for 62%, 77% and 79% of ID-A variance in ALS, FTD and AD, respectively (Figure 5D), strongly supporting that ID-A Indels lead to DNA fragmentation in a disease-specific process. Interestingly, we also observed significantly increased levels of pre-existing double-strand breaks in diseased brains, which similarly correlated with ID-A burdens (Figures 5A–5D), suggesting that progressive accumulation of single-strand breaks may eventually lead to double-strand break formation and widespread genome instability in disease.

**Figure 5.**
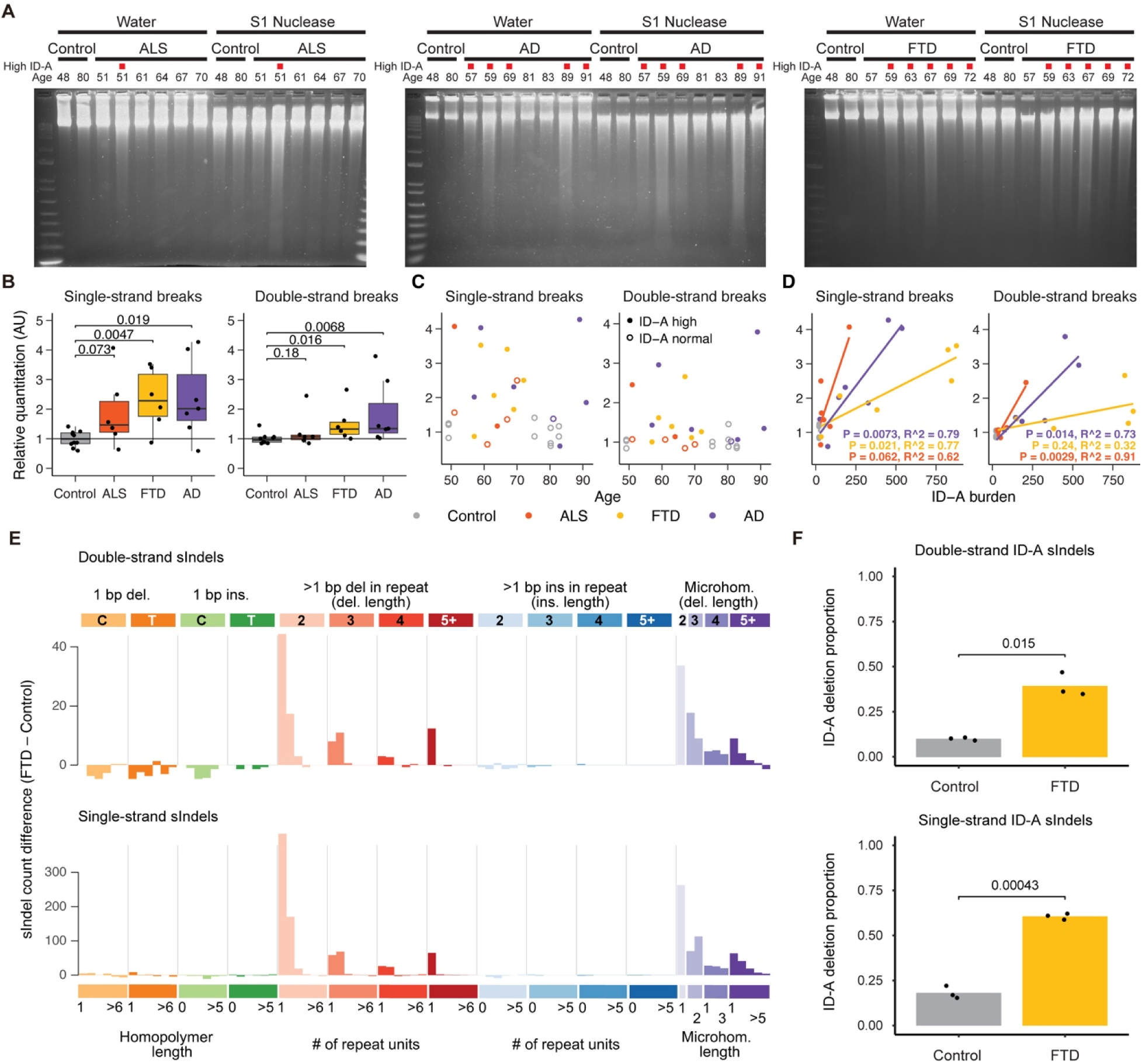
Double-strand ID-A sIndels associate with single-strand breaks and deletions. (A) Agarose gel electrophoresis of gDNA from motor and premotor cortex tissues of *C9ORF72* ALS brains and premotor cortex tissues of control brains, as well as from prefrontal cortex tissues of *C9ORF72* FTD, AD and control brains. Samples are arranged in order of donor age. Increased smear intensity following S1 nuclease treatment indicates single-strand breaks. Red squares indicate ID-A-high cases determined from single cell WGS (see STAR Methods) and strongly correlate with the appearance of increased smear intensity following S1 nuclease treatment on the gel. (B) Mean single-strand break and double-strand break intensities for each condition (*C9ORF72* ALS, *C9ORF72* FTD, AD, and control). *P*-values are Wilcoxon rank-sum tests. Y-axis values represent quantification of the smears in (A) in arbitrary units (AU) using Agilent TapeStation. (C) Abundance of single-strand and double-strand break intensities from panel (B) plotted against age of donor. (D) Average ID-A burden per brain is correlated with single-strand break and double-strand break intensities quantified in (B) in *C9ORF72* ALS, *C9ORF72* FTD and AD brains, suggesting a common process. Each point represents one brain; averages are taken over all cortical neurons subjected to scWGS from each brain. *P*-values are regression *t*-tests on slope. *P*-values for controls are not significant and omitted on the graph. (E) Indel spectra derived from duplex sequencing of 50-neuron mini-bulk samples with META-CS. Spectra are shown by subtracting the mean spectrum of age-matched neurotypical controls from the mean spectrum of *C9ORF72* FTD cases. Spectra for double-strand and single-strand sIndels, which can be distinguished due to strand-specific tagging, are shown separately. (F) FTD shows increased proportions of ID-A-like deletions among both single- and double-strand sIndels (numbers of ID-A-like deletions divided by total numbers of Indels) compared to age-matched controls from META-CS duplex sequencing data. *P*-values are two-tailed unpaired *t*-tests.

Multiplexed end-tagging amplification of complementary strands (META-CS)^49^, a type of duplex sequencing which distinguishes the Watson and Crick strands of DNA molecules, provided a third, orthogonal confirmation of the presence of single- and double-strand 2-bp deletions in diseased brains. During TOP1-mediated RER, the subsequent re-ligation of single-strand breaks creates an intermediate DNA product containing a 2-bp single-strand deletion (Figure 4A). We performed META-CS on 50-neuron samples from the prefrontal cortex from three ID-A-high *C9ORF72* FTD brains and three age-matched control brains. The duplex sequencing data confirmed that ID-A-like, double-strand 2-bp deletions were enriched in *C9ORF72* FTD brains, as evidenced by the same deletion being captured on the independently-sequenced Watson and Crick strands. Duplex sequencing also captured a larger number of ID-A-like single-strand lesions in *C9ORF72* FTD brains, for which the deletion was absent on one of the two independently sequenced strands (Figure 5E). The proportions of ID-A-like deletions were significantly higher in *C9ORF72* FTD brains compared to control brains for both double-and single-strand events, but single-strand deletions showed a more prominent increase (Figure 5F).

These results suggest that the increased levels of TOP1-mediated RER in neurons from neurodegenerative conditions generate frequent single-strand breaks and deletions, a small fraction of which are misrepaired to create double-strand deletions. Consistent with this, diseased neurons show upregulated TOP1 expression but reduced expression of pathways responsible for clearing TOP1ccs (Figure S8). Together, these findings indicate that increased TOP1-mediated RER is a common mechanism underlying widespread, catastrophic DNA damage in neurodegenerative conditions.

### TOP1-mediated mutagenesis is absent in the cerebellum of diseased brains

To determine whether the increased TOP1-induced DNA damage and somatic mutations observed in diseased neurons reflect region-specific pathological processes rather than nonspecific end-stage genomic DNA damage, we analyzed somatic mutations and DNA strand breaks in the cerebellum, a brain region that is relatively spared in ALS, FTD, and AD. We performed PTA-based scWGA and scWGS on three cerebellar granule cell neurons from each of the diseased brains for which cortical neurons were collected (Figures 6A and S9)^50,51^. Analysis of scWGS data revealed an absence of 2-bp deletions in the sIndel spectra of cerebellar neurons across all three neurodegenerative conditions (Figure 6B). Consistent with this, the fractions of 2-bp deletions were significantly lower in cerebellar neurons compared to cortical neurons (Figure 6C). The overall burdens of ID-A sIndels were also significantly lower in cerebellar neurons compared to cortical neurons (Figure 6D). Stratification of cerebellar neurons by ID-A status further demonstrated that only a single cerebellar neuron from a *C9ORF72* ALS brain met criteria for ID-A–high classification, whereas no ID-A–high cerebellar neurons were detected in *C9ORF72* FTD or AD cases (Figure 6E).

**Figure 6.**
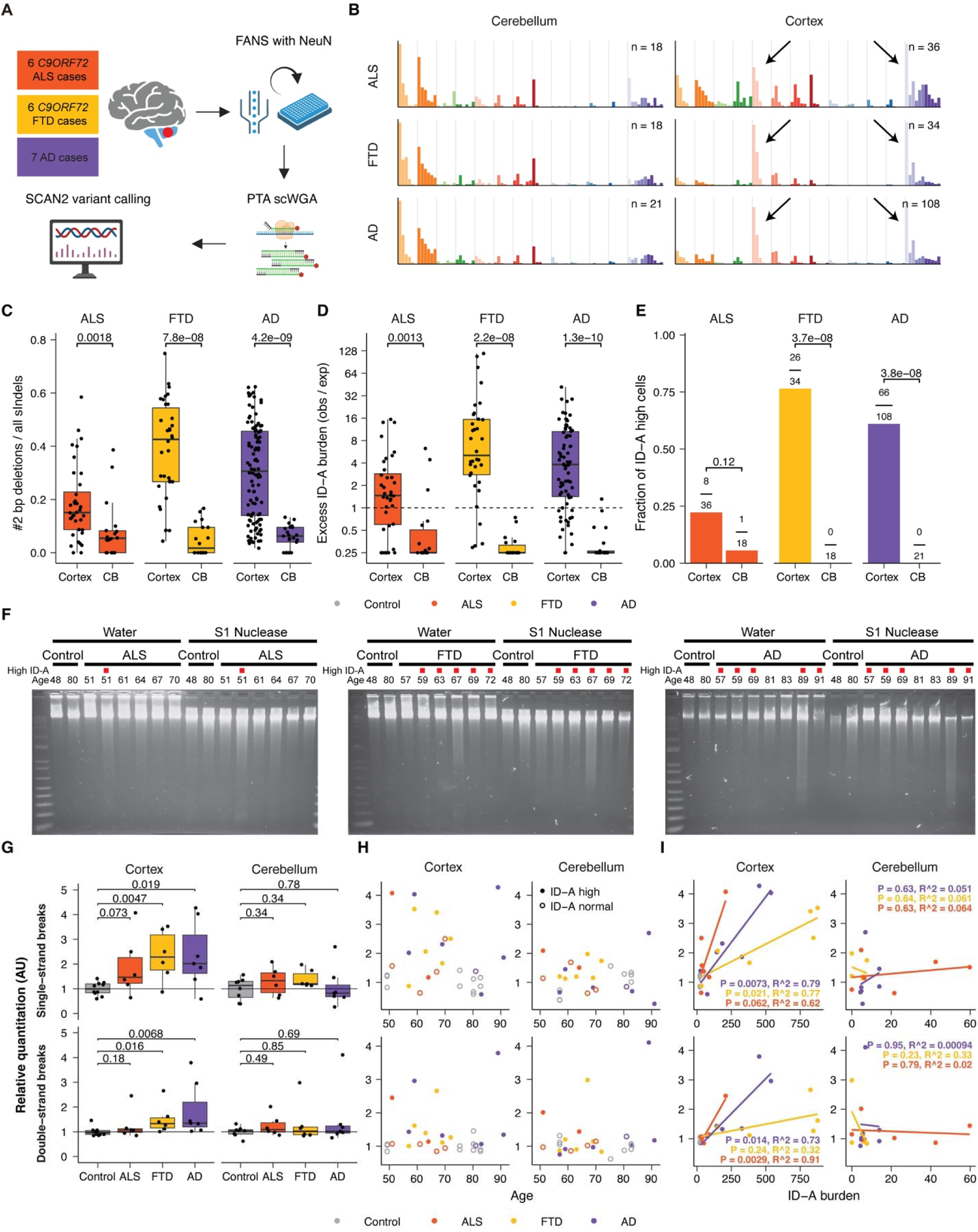
Cerebellar samples from *C9ORF72* ALS, *C9ORF72* FTD and AD brains do not show evidence of TOP1-mediated mutagenesis. (A) Overview of the experimental design. Three cerebellar granule neuron nuclei were isolated from the cerebellum of each of the 6 *C9ORF72* ALS, 6 *C9ORF72* FTD, and 7 AD brains for which scWGS of cortical neurons were analyzed in this study. Cerebellar neurons were then amplified by PTA, subjected to genome sequencing, and analyzed using the same procedures as cortical neurons. (B) sIndel spectra of cerebellar neurons and cortical neurons from diseased brains. (C) Fraction of 2 bp deletions in cortical and cerebellar neurons from diseased brains. *P*-values represent Wilcoxon rank-sum tests. CB: cerebellum. (D) Excess burdens of ID-A sIndels in cortical and cerebellar neurons from diseased brains shown as observed/expected ratios. Expected values were derived from PTA-amplified control cortical neurons. *P*-values represent Wilcoxon rank-sum tests. (E) Fraction of cells with high levels of ID-A sIndels in cortical and cerebellar neurons from diseased brains, determined by a classifier (see STAR Methods); *P*-values represent Fisher tests of independence between disease status and ID-A-high classifications. (F) Agarose gel electrophoresis of gDNA from cerebellum tissues of disease and control brains. Samples are arranged in order of donor age. Increased smear intensity following S1 nuclease treatment indicates single-strand breaks. Red squares indicate ID-A-high cases determined from single cell WGS (see STAR Methods). (G) Mean single-strand break and double-strand break intensities in cortex and cerebellum for each condition (*C9ORF72* ALS, *C9ORF72* FTD, AD, and control). *P*-values are Wilcoxon rank-sum tests. Y-axis values represent quantification of the smears in (F) in arbitrary units (AU) using Agilent TapeStation. (H) Abundance of single-strand and double-strand breaks in cortex and cerebellum vs. age. (I) ID-A burden from single-cell WGS of cerebellar neurons (mean observed/expected ratio across neurons in each case) does not correlate with single-strand break and double-strand break intensities quantified in (F) for *C9ORF72* ALS, *C9ORF72* FTD and AD brains. *P*-values are regression *t*-tests on slope. *P*-values for controls are not significant and omitted on the graph. See also Figure S9

We next performed strand break analyses on gDNA extracted from cerebellum tissues of diseased and control brains. In contrast to cortical neurons, where diseased brains showed elevated strand-break levels, levels of both single-strand breaks and double-strand breaks in cerebellar gDNA were comparable between diseased and control brains (Figures 6F and 6G) and did not correlate with donor age or ID-A burdens in cerebellar neurons (Figures 6H and 6I). Together, these findings indicate that TOP1-mediated mutagenesis is largely absent in the cerebellum of diseased brains and is restricted to disease-relevant cortical regions, consistent with a region-specific contribution of TOP1-induced genomic damage to ALS, FTD, and AD pathogenesis.

### Deficiency of RNase H2 and ribonucleotide reductase in diseased neurons

During DNA repair, DNA polymerases must incorporate dNTPs and discriminate against rNTPs. This selectivity relies on polymerase properties, as well as the balance of dNTP and rNTP pools, which is regulated by ribonucleotide reductases (RNRs)^52^: when the dNTP:rNTP ratio is low, rNTP incorporation increases, threatening genome stability. Embedded rNTPs are removed by RNase H2, but in its absence, TOP1 serves as an alternative repair pathway (Figure 7A). We found that genes encoding RNRs and RNase H2 subunits were downregulated across various neuronal subtypes in our snRNA-seq data from *C9ORF72* FTD brains compared to controls (Figure S10A). A similar trend of reduced expression was also observed in previously published snRNA-seq data from AD neurons (Figure S10A), although most of these changes were not statistically significant, likely due to the low abundance of these transcripts in neurons and differences in tissue lysis and nuclear isolation protocols. However, qPCR on cDNA from neurons of *C9ORF72* FTD and AD brains with high levels of ID-A sIndels and control brains confirmed significantly reduced expression of *RNASEH2B* and *RRM2B* in *C9ORF72* FTD and AD neurons compared to control neurons (Figures 7B and S10B).

**Figure 7.**
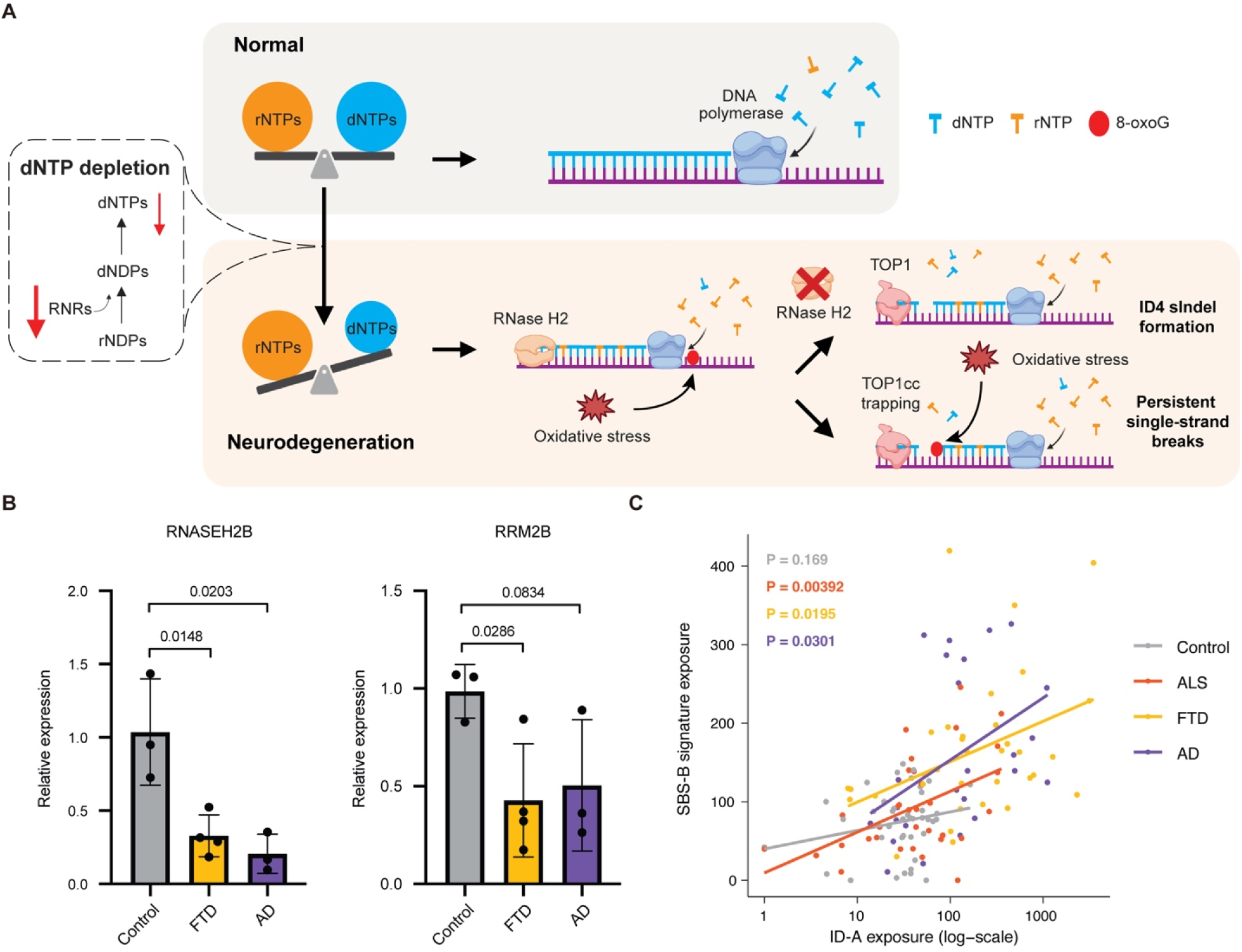
Potential mechanisms underlying the dysregulation of RER. (A) Model of TOP1-mediated mutagenesis in neurons from neurodegenerative conditions. RNR: ribonucleotide reductase. (B) Expression levels of *RNASEH2B* and *RRM2B* in neurons measured by qPCR. *P*-values denote two-tailed unpaired *t*-tests. (C) Correlation between burdens of *de novo* signatures SBS-B (sSNVs) and ID-A (sIndels) in PTA-amplified cortical neurons. SBS-B resembles the COSMIC SBS30 signature, which is associated with a deficiency in NTHL1 and base excision repair of oxidative DNA damage repair. *P*-values indicate regression *t*-tests on slope. See also Figure S10 and S11

Finally, we observed a strong correlation between ID-A and SBS-B—which resembles COSMIC SBS30, a signature associated with oxidative DNA damage—in cortical neurons from all three disease conditions after removing three outlier ALS neurons that exhibited elevated sSNV burdens, possibly driven by another mechanism not involving TOP1 (Figures 7C, S10C, and S11). The correlation between elevated sSNV and sIndels in the same neurons indicates that neurons that accumulate higher levels of DNA damage due to oxidized nucleotides are the same ones that show catastrophic increases in TOP1-mediated mutagenesis. This correlation could reflect the known tendency of DNA polymerases β and λ to erroneously incorporate ribonucleotides into the genome during excision repair of oxidative DNA damage^53^.The misincorporation of ribonucleotides under these conditions could exacerbate genomic instability, further amplifying TOP1-mediated mutagenesis (Figure 7A). Furthermore, this correlation between oxidative DNA damage and ID-A mutagenesis may contribute to the increased single-strand breaks observed in diseased brains, as local oxidative DNA damage can block the re-ligation of single-strand breaks by trapping TOP1cc, resulting in persistent single-strand breaks (Figure 7A)^54–56^.

## DISCUSSION

Our results revealed an accelerated accumulation of both sSNVs and sIndels in neurons from *C9ORF72* ALS, *C9ORF72* FTD and AD brains, and further suggest that these processes are correlated and central to the pathogenic process of these conditions with highly divergent pathological appearance. While sIndels accumulate at a much slower rate than sSNVs in normal neurons—reaching approximately 200 to 300 sIndels in elderly individuals^26^— this process is greatly accelerated in neurons affected by these three neurodegenerative conditions, with some neurons exhibiting several thousand sIndels (Figure 2A), equivalent to hundreds of years of age-related sIndel accumulation. Intriguingly, the excessive sIndels in neurons from all three diseases shared many key features that strongly resemble the COSMIC ID4 signature (Figures 3C and 3F). COSMIC ID4 has been linked to RNase H2 deficiency and alternative TOP1-mediated RER^41^ and is also the most prevalent aging-associated Indel signature in normal neurons^26^. Consistent with this, age-related downregulation of *RNASEH2C* was detected across multiple neuronal cell types in normal human brains, but not in glial cells, potentially predisposing neurons to neurodegeneration^57^. Furthermore, we observed several characteristics of TOP1-mediated deletions, including the local TNT motif, the most commonly deleted dinucleotide (CT), the direct nuclear enrichment of TOP1cc immunoreactivity and increased TOP1cc in disease neurons, and the association with local gene expression level. In addition, we observed other expected side effects of TOP1 activity, such as a preponderance of single-strand breaks and precursor single-strand deletions. Taken together, our evidence strongly suggests a role for TOP1 in somatic mutagenesis in neurodegenerative conditions of diverse phenotypes and pathologies.

TOP1 is a DNA topoisomerase that facilitates transcription, DNA replication, and DNA repair by alleviating topological tensions in the DNA^55,56,58^. During this process, TOP1 forms a transient cleavage complex (TOP1cc), in which it covalently binds to DNA and introduces a single-strand nick that is subsequently re-ligated. In neurons, TOP1 has been implicated in the transcriptional regulation of long neuronal genes essential for synaptic function and plasticity^59,60^, and depletion of TOP1 in neurons has been linked to early-onset neurodegeneration and autism^59,61^. In the absence of RNase H2, TOP1 compensates by facilitating the removal of misincorporated ribonucleotides from the genome. Single-strand breaks induced by TOP1 can produce intermediate DNA products with single-strand 2-bp deletions, which were detected by our duplex sequencing approach (Figures 5E and 5F). Given the increased oxidative damage observed in neurons from the three neurodegenerative conditions manifested as increased oxidative sSNV signature exposure (Figures S4A and S4B, SBS-B), the increased level of single-strand breaks in *C9ORF72* FTD and AD brains is likely a result of TOP1cc trapping by local oxidative DNA damage (Figures 5A-5D). Thus, although sSNVs caused by oxidative stress may not be sufficiently numerous to affect the genome, the resulting catastrophic levels of somatic deletions present in some neurons could potentially contribute to neuronal dysfunction by impacting the transcription of hundreds or even thousands of genes. Furthermore, the strong correlation between ID-A deletions and single-strand breaks (Figures 5A-D) suggests that DNA fragmentation—a hallmark feature of these neurodegenerative conditions^9,11^—is not a result of poor tissue quality or other technical factors; instead, it occurs during the neuron’s lifetime as a consequence of TOP1 activity during RER and may be relevant to disease progression. In addition to oxidative DNA damage-induced TOP1cc trapping, impaired repair of trapped TOP1ccs may further contribute to their persistence. Our snRNA-seq analyses showed that genes involved in TOP1cc resolution are downregulated (Supplementary Fig. S8). These include components of the SPRTN-mediated proteolysis and TEX264-dependent nucleophagy pathways^62,63^, both of which are VCP/p97-dependent TOP1cc repair pathways. Importantly, VCP is a gene mutated in some familial ALS and FTD patients. Together, these observations suggest that impaired TOP1cc clearance may further exacerbate the accumulation of DNA damage and genomic fragmentation in vulnerable neurons.

The observation of a shared mutagenic mechanism between such categorically different conditions—TDP-43 (ALS and FTD) and tau proteinopathies (AD)—is both surprising and suggestive of even broader applicability to neurodegenerative disease and perhaps neuronal death in general. Indeed, a recent report identified ID-A-like deletions in chronic traumatic encephalopathy (CTE)^64^, a second tauopathy caused by repetitive head trauma^65^. TDP-43 pathology has been observed in a subset of AD and CTE cases^66,67^, raising the possibility that nuclear TDP-43 depletion is the underlying shared cause of ID-A deletions; however, this appears unlikely since we did not observe an association between ID-A deletions and TDP-43 depletion status in *C9ORF72* ALS and FTD brains (Figures 2D and S6), and TDP-43 knockdown in iPSC-derived neurons did not induce TOP1cc formation (Figure S12). Importantly, we also excluded the possibility that tissue preservation artifacts contribute to the observed mutational pattern. Brain samples were obtained from multiple independent sources (Table S1)^64^, and levels of somatic mutations, particularly ID-A deletions, did not correlate with postmortem interval across samples (Figure S13). Together, these results indicate that neither TDP-43 depletion nor tissue quality accounts for the shared ID-A mutational signature. Instead, increased oxidative stress—implicated in all of these neurodegenerative disease conditions^7,10,68^—may play a key role. The strong correlation between ID-A deletions and oxidative sSNV signature exposure (Figure 7C) supports oxidative stress as a plausible driver of ID-A deletion formation across a wide range of neurodegenerative conditions. Indeed, oxidative DNA damage can increase the erroneous incorporation of ribonucleotides into the genome during excision repair of oxidative DNA damage and elevate the burden of substrates requiring RNase H2-dependent RER. In addition, *RNASEH2A* expression is reduced in Werner syndrome, a progeroid disorder characterized by replication stress and DNA damage^69^, suggesting that genotoxic stress can suppress RNase H2 expression. Therefore, oxidative DNA damage may both increase ribonucleotide incorporation and secondarily reduce RNase H2 expression, thereby further fueling TOP1-dependent RER activity. One plausible upstream cause of the oxidative stress itself would be the inflammatory activation of microglial cells, which has been observed in all of these conditions^70^, and which is associated with increased secretion of cytokines and reactive oxygen species that may damage nearby neurons.

The shared genomic pathology in diverse neurodegenerative conditions like ALS, FTD and AD raises the question of how general this pattern of genomic damage is to other forms of neuronal degeneration. Since our ALS and FTD cohorts all harbored *C9ORF72* repeat expansions, our findings may not necessarily generalize to all ALS and FTD cases, especially those without known genetic mutations. Nonetheless, diseases such as Ataxia telangiectasia and Aicardi-Goutières syndrome likely also involve ID-A deletions, as these conditions involve persistent TOP1cc or RNase H2-related RER deficiency^71,72^, which our data suggests is predictive of ID-A deletions. Since our ALS and FTD cases involved *C9ORF72* repeat expansions, it is also reasonable to question whether other repeat expansion disorders would harbor ID-A deletions. Somatic expansion of unstable repeats, such as those in trinucleotide repeat disorders, often involves activation of various DNA damage repair pathways, which may deplete the cellular dNTP pools and exacerbate ribonucleotide incorporation into the genome^73^. Overall, our data support that preventing excessive or unresolved TOP1cc formation and/or improving RER activity—rather than direct TOP1 inhibition—may represent potential therapeutic targets in a broad spectrum of neurodegenerative conditions.

## RESOURCE AVAILABILITY

### Lead contact

Requests for further information and resources should be directed to and will be fulfilled by the lead contact, Christopher A. Walsh.

### Materials availability

This study did not generate new unique reagents.

### Data and code availability

Newly generated single-cell DNA sequencing data from neurotypical controls have been uploaded to dbGaP accession ID phs001485.v3.p1. Bulk and single-cell DNA- and RNA-sequencing data from ALS and FTD cases have been deposited at the National Institutes on Aging’s data depository NIAGADS. All newly generated sequencing data are available conditional on approval for controlled access, in line with privacy regulations. Previously published bulk whole genome and single-cell DNA-, RNA- and ATAC-sequencing data are available for download at dbGaP, accession ID phs001485.v3.p1, and NIAGADS, accession NG00162, subject to controlled data access approval. All scripts used to perform analyses are available on Zenodo at https://doi.org/10.5281/zenodo.20174164.

## ACKNOWLEDGMENTS

We thank the Massachusetts Alzheimer’s Disease Research Center and NIH NeuroBioBank for providing fresh frozen human tissues. We thank the donors and families for their contributions, and J. E. Neil for assistance with tissue procurement. We thank the Research Computing group at Harvard Medical School and Boston Children’s Hospital. We thank Dr. Paul McKeever for discussion on snRNA-seq data. dCAS9-ZIM3-KRAB-BFP construct was a kind gift from Dr. Michael E. Ward at National Institute of Health. Dual guide RNA lentiviral constructs were a kind gift from Dr. Paul C. Blainey at Broad Institute of MIT and Harvard^74^. This work was supported by the PRMRP Discovery Award W81XWH2010028 (Z.Z.); the Edward R. and Anne G. Lefler Center postdoctoral fellowship (Z.Z.); the American Heart Association Career Development Award 23CDA1046074 (Z.Z.); Fred and Gilda Slifka Neuroscience Transformative Scholar Award (N.R.); DP2 AG086138 (M.B.M.); R01 AG082346 (M.B.M.); R56 AG079857 (A.Y.H.); R01 AG088082 (A.Y.H.); the Alzheimer’s Association Research Fellowship (A.Y.H.); R01 HG012573 (P.J.P.); RM1 NS133601 (C.L.-T.); K01 AG051791 (E.A.L.); the Suh Kyungbae Foundation (E.A.L.), DP2 AG072437 (E.A.L.); R01 NS032457 (C.A.W.); R01 AG070921 (C.A.W. and E.A.L.); UG3 NS132138 (C.A.W. and P.J.P.) and UG3 NS132144 (E.A.L. and C.A.W.) through the SMAHT Consortium; a Massachusetts Alzheimer’s Disease Research Center pilot grant (C.L.-T. and C.A.W.); and the Allen Family Philanthropies (C.A.W. and E. A. L.). C.L.-T. is supported by the Araminta Broch-Healey Endowed Chair in ALS. C.A.W. is an Investigator of the Howard Hughes Medical Institute

## AUTHOR CONTRIBUTIONS

Conceptualization, Z.Z., L.J.L., G.D., P.J.P., C.L.-T., E.A.L., C.A.W.; Methodology, Z.Z., L.J.L., G.D., D.D.S., W.J.N., A.N.; Resource, K.E., E.G., M.B.M., A.Y.H., C.L.-T, Software, L.J.L., G.D.; Investigation, Z.Z., L.J.L., G.D., Junho K., Jayoung K., K.K., N.R., M.B., A.C., B.S.; Visualization, Z.Z., L.J.L., G.D., Jayoung K.; Funding acquisition, Z.Z., P.J.P., C.L.-T., E.A.L., C.A.W.; Project administration, Z.Z., L.J.L., P.J.P., C.L.-T., E.A.L., C.A.W.; Supervision, Z.Z., L.J.L., P.J.P., C.L.-T., E.A.L., C.A.W.; Writing – original draft, Z.Z., L.J.L., G.D.; Writing – review & editing: Z.Z., L.J.L., G.D., Junho K., D.D.S., B.S., M.B.M., A.Y.H., P.J.P., C.L.-T., E.A.L., C.A.W.

## STAR METHODS

### EXPERIMENTAL MODEL AND STUDY PARTICIPANT DETAILS

#### Human brain tissue sources

Fresh frozen postmortem human brain tissues were originally collected by the Massachusetts Alzheimer’s Disease Research Center and NIH NeuroBioBank (Table S1) according to their respective institutional protocols, written authorization and informed consent; they were subsequently obtained for this study with the approval of the Boston Children’s Hospital Institutional Review Board. Research on these deidentified specimens and data was performed at Boston Children’s Hospital with approval from the Committee on Clinical Investigation. The presence of *C9ORF72* repeat expansion in these cases were determined through repeat-primed PCR (RP-PCR).

### METHOD DETAILS

#### Neuronal nuclei isolation

Single neuronal nuclei were isolated using FANS with co-staining of NeuN and TDP-43 antibodies^27,75^. Fresh frozen postmortem human brain tissues were homogenized in a dounce homogenizer with a chilled tissue lysis buffer (10mM Tris-HCl, 0.32M sucrose, 3mM Mg(OAc)2, 5mM CaCl2, 0.1mM EDTA, 1mM DTT, 0.1% Triton X-100, pH 8) on ice. Tissue lysates were carefully overlayed on top of a sucrose cushion buffer (1.8M sucrose 3mM Mg(OAc)2, 10mM Tris-HCl, 1mM DTT, pH 8) and ultra-centrifuged for 1 hour at 30,000 x g. Nuclear pellets were incubated and resuspended in ice-cold PBS supplemented with 3mM MgCl2, filtered (40 μm pore size), then stained with Alexa Fluor 488-conjugated anti-NeuN antibody (Millipore MAB377X) and Alexa Fluor 647-conjugated anti-TDP-43 antibody (Abnova H00023435-M01). DAPI was added to stained nuclei right before the sorting. Single neuronal nuclei with TDP-43 pathology (NeuN+, TDP-43-) and without TDP-43 pathology (NeuN+, TDP-43+) were then sorted into individual wells of 96-well plates

#### Single-cell whole-genome amplification

Primary template-directed amplification (PTA) of sorted single nuclei was performed using the ResolveDNA® Whole Genome Amplification Kit v1 (BioSkryb 100136) according to the manufacturer’s protocol^26^. Amplified single-cell genomes were subjected to quality control using Picogreen quantification and a 4-locus multiplex PCR assay to assess DNA yield and amplification evenness^75^. Single-cell genomes with sufficient DNA yield and successful amplification of all four loci were selected for single-cell whole-genome sequencing (scWGS).

#### Library preparation for scWGS

Libraries were made from PTA amplified single-cell genomes following a modified KAPA HyperPlus Library Preparation protocol provided by BioSkryb ^26^. Libraries were subjected to quality control using Picogreen quantification and Tapestation HS D1000 Screen Tape (Agilent PN 5067–5584) before sequencing. Single cell genome libraries were sequenced on the Illumina NovaSeq platform (150 bp x 2) at 30X for cortical neurons and 22X for cerebellar neurons.

#### scWGS data preprocessing

Illumina reads were aligned to the human reference with decoy sequence GRCh37d5 (hs37d5) using bwa mem version 0.7.15-r1140 with the -M option. Following alignment, BAMs were processed according to the GATK Best Practices recommendations, namely Picard MarkDuplicates version 2.8.0 and IndelRealigner, BaseRecalibrator and PrintReads using GATK version 3.5.0-g36282e4.

#### scWGS somatic mutation calling with SCAN2

Somatic SNVs and Indels were called from PTA and MDA data using SCAN2 version 1.2.13, git commit `09ed5b9` and r-scan2 git commit `c0aef34`. The standard SCAN2 recommended workflow was followed. First, to enable somatic Indel calling, `scan2 makepanel` was run on 8 separate batches of of PTA and MDA single cells and matched bulks to make the analysis tractable. The samples included in each batch are defined in the scripts provided at https://doi.org/10.5281/zenodo.20174164. Each batch included the 56 control PTA cortical neurons and matched bulks generated in Ganz et al.^35^ to identify recurrent indel artifacts with at least the same specificity as previous published analyses. Second, one instance of `scan2 call_mutations` was run for each unique individual using the makepanel output corresponding to that individual’s batch. For each run, all BAMs belonging to the individual (i.e., all MDA, PTA and matched bulk(s)) were provided to SCAN2. Third, one instance of SCAN2’s mutation-signature based rescue (`scan2 rescue`) was run for each unique combination of amplification type (PTA or MDA), phenotype (ALS, FTD, AD or control) and cell type (neuron, oligodendrocyte or mixed glia). Finally, SCAN2’s post-processing script to filter clustered or recurrent (i.e., called in >1 single cell) artifacts and mutations (`SCAN2/bin/digest_calls.R`) was applied to each `scan2 rescue` output. For mutation enrichment analysis, only calls with `final.filter=FALSE` were retained.

Critically, because of the unique mutational signature of disease-associated Indels, we opted to exclude all SCAN2 rescue mutation calls from all analyses to avoid any bias involved in signature-based rescue. Rescued calls were removed by excluding rows with `rescue=TRUE` from the final recurrence filtered output. Additional configuration options applicable to all SCAN2 steps were `--gatk=sentieon_joint`; the GRCh37 human reference genome with decoy hs37d5 (`--ref`), dbSNP v147 common (`--dbsnp`) and 1000 Genomes phase 3 SHAPEIT2 phasing panel (`--shapeit-refpanel`).

#### Single-cell QC and SCAN2 performance metrics

Technical metrics (mean sequencing depth, MAPD) and mutation detection metrics (sensitivity, fraction of genome analyzable and false discovery rates) were obtained using SCAN2’s `callstats()` function in `R`. The median absolute pairwise difference (MAPD) values were extracted from each single cell using SCAN2’s `mapd()` function. MAPD measures non-uniformity of amplification by examining the difference in read count between neighboring genomic bins: MAPD = median_i_ {|log_2_ R_i_ - log_2_ R_i+1_|} where Ri is the GC-corrected read depth in genomic bin *i* divided by the average GC-corrected read depth across the genome. The set of genomic bins is distributed with the SCAN2 package and aims to create bins with an equal number of alignable bases using paired end 150 bp Illumina reads. MAPDs reported here used a bin size of 64 kb. SCAN2 estimates somatic SNV and Indel detection sensitivity by applying somatic criteria to known germline variants detected in the matched bulk^26^. The fraction of genome analyzable for each single cell is the number of positions in the reference genome hs37d5 that meet the minimum sequencing read depth in the matched bulk (>10 reads) and single cell (>5 reads for sSNVs, >9 reads for sIndels) divided by the number of positions accessible by short read sequencing, estimated by the number of positions with >0 reads in the matched bulk. False discovery rates (FDRs) were derived from previous estimates of false positive errors committed by SCAN2 per megabase of genome analyzed that relied on *in silico* benchmarking experiments that preserved single-cell artifacts^26^. Briefly, read data from haploid X chromosomes from two male single cells were combined to create a synthetic diploid X chromosome, which was then analyzed by SCAN2. True somatic mutations—which should have allele fractions of ∼100% on haploid chromosomes—were identified prior to combining the read data. Any SCAN2 mutation call not identified as a true mutation was counted as a false positive. Pseudoautosomal regions were excluded from this analysis. This process was performed for several pairs of male single cells to arrive at a false positive rate of 0.013 sSNV errors per Mb and 0.00073 sIndel errors per Mb. For each single cell and mutation type (SNV or indel), the false positive load is the product of multiplying the respective rate by the respective number of analyzable bases in that cell. The per-cell FDR is the false positive load divided by the total number of mutation calls or 1, whichever is smaller. The catalog-wide FDR is the sum of false positive loads over all cells divided by the sum of calls over all cells.

#### scWGS mutation burden estimation and comparison

SCAN2 estimates the genome-wide mutation burden by correcting the number of called mutations for detection sensitivity. This correction accounts for sensitivity changes caused by sequencing depth, imbalanced and/or non-uniform single-cell amplification, and other factors. Correction is performed separately for sSNVs and sIndels. More details are provided in our previous publication^26^. SCAN2’s sSNV and sIndel mutation burden estimates were extracted from each SCAN2 object in R using r-scan2’s `mutburden()` function. To compare mutational burdens between any two groups (e.g., control and ALS), it is first necessary to correct for differences in age between the two groups. Two strategies were used together to achieve this goal. First, we computed the expected burden as a function of age using PTA-amplified, neurotypical control neurons. This process involved two steps of linear models; the first to identify strong outliers and the second to compute the final age-based expectation. In more detail, the first linear model was fit to all PTA control neurons via `lm(burden ∼ age)` and Tukey’s method was applied to the model’s residuals to identify strong outliers. The second linear model was then fit with the outliers removed and R’s `predict` function was used to derive the expected burden as a function of age. Separate models were fit to sSNV, sIndel and mutational signature burdens (see *Mutational signature analysis* for more details on signature burdens); for analysis of oligodendrocytes, n=66 PTA-amplified control oligodendrocytes were used in place of PTA control neurons. For each cell and burden type (i.e., sSNV, sIndel, etc.) a final observed/expected ratio was produced by dividing the observed burden from SCAN2 by the expected burden based only on the cell’s age from the linear model. Second, for all burden comparisons between groups, only observed/expected ratios from cells with age > 50 were included since the residuals of younger cells tended to be large compared to the trend line, leading to inflated observed/expected ratios. Significance tests for differences in burdens between groups are Wilcoxon rank-sum tests on the observed/expected ratios. After acceptance of the manuscript, we identified that control case UMB5451 had been incorrectly annotated as female instead of male. Reanalysis using the corrected sex annotation showed no meaningful changes in mutation calls or reported results. Because SCAN2 performs mutation calling independently of sample sex and the analyses presented in this study report only autosomal mutation burdens, the incorrect sex annotation did not materially affect the conclusions of the study. Therefore, the figures and tables in the manuscript were not modified.

#### scWGS quality control and removal of outliers

To ensure that technical factors did not drive differences in burdens or mutational signatures, we compared sSNV and sIndel burdens against a quantity that reflects quality of single cell amplification, the median absolute pairwise difference (MAPD)^36^. Low MAPD values indicate more uniform amplification and thus better quality. Plots of mutation burden vs. MAPD were visually assessed to identify cells in which technical quality may have driven high sIndel burdens (i.e., cells with both high sIndel and high MAPD). This revealed 5 PTA-amplified neurons (3 AD, 2 FTD) and 7 MDA-amplified neurons (6 AD, 1 control), which were excluded from further analysis. Additionally, 10 MDA-amplified neurons were excluded due to extreme sIndel burden and/or a highly atypical mutation signature enriched in long microhomology deletions.

#### Mutational signature analysis

Matrices of SBS96 and ID83 somatic mutation counts were generated with SigProfilerMatrixGenerator version 1.2.26 using the GRCh37 human reference genome^76^. All single cells (both MDA- and PTA-amplified; Table S2), including outliers, were included for matrix generation. Next, sIndel count matrices were corrected to reflect different sensitivities for the 83 Indel channels, as estimated in our previous publication^26^. De novo mutational signatures were then extracted using SigProfilerExtractor version 1.1.24 and options min_stability=0.9, minimum_signatures=1, maximum_signatures=8, and nmf_replicates=100^77^. The decompositions with the most stable signatures were selected, which was K=6 signatures for SBS96 and K=3 signatures for ID83. For each de novo signature, a genome-wide burden was computed by multiplying the SigProfilerExtractor assigned number of mutations by SCAN2’s genome-wide scaling factor for the appropriate mutation type (i.e., SBS96 counts were multiplied by SCAN2’s sSNV genome-wide burden scaling factor and ID83 counts were multiplied by SCAN2’s sIndel genome-wide scaling factor). These scaled genome-wide burdens were then used to fit models on PTA-amplified control neurons and calculate age-adjusted expected values using the procedure detailed in *scWGS mutation burden estimation and comparison*.

#### Classification of ID-A high and normal cells

To label each single cell either ID-A normal (typical levels of ID-A) or ID-A high (excessive ID-A, consistent with disease progression), we built a classifier based on 83-dimensional sIndel spectra. A matrix of sIndel spectra was first constructed using the same cells and corrected indel spectra described in *Mutational signature analysis*. We then used the R `umap` package to perform dimensional reduction from the 83-dimensional spectra into 2-dimensional space via the function call `umap(M, preserve.seed=FALSE, random_state=1, transform_state=1)`, where M represents the matrix of indel spectra. Under this dimensional reduction, all cells visually clustered into two distinct populations, one of which corresponded to high levels of ID-A. Cells in the cluster with elevated ID-A levels were classified as ID-A high; all other cells were classified as ID-A normal. Cells with no sIndel calls (n=2 PTA-amplified control cortical neurons and n=3 PTA-amplified control oligodendrocytes) were not classified.

#### Mutation enrichment analysis

*Enrichment tests.* Enrichment analysis was performed following the procedures outlined in our previous publications^26,35^. Briefly, enrichment analysis considers the number of somatic mutations over a set of genomic regions which are not, in general, contiguous (e.g., defined by covariates such as gene expression). The null hypothesis is that somatic mutations are uniformly distributed over the genome. To test the null hypothesis, SCAN2 simulates uniformly distributed mutations across the genome for each single cell, considering both (1) the regions of the genome with sufficient read depth to detect somatic mutations and (2) the mutational spectrum of the whole mutation set, which is required to match the observed somatic mutations. This simulation is performed 10,000 times to produce 10,000 null sets of mutations. The true mutation sets and null sets of mutations are then combined for each set of single cells of interest (e.g., all PTA single neurons from ALS). For each genomic region, the true mutation set and simulation sets are intersected with the region and tallied. The enrichment over a region is defined as the number of true mutations in the region divided by the mean number of simulated mutations in the region over the 10,000 simulations; values > 1 indicate enrichment while values < 1 indicate depletion. This same metric is be applied to all 10,000 simulations to construct a distribution of the enrichment statistic. A two-sided enrichment test *P*-value is constructed by counting the number of simulations with more extreme enrichment ratios than the observed mutation count (to make this test two-sided, more extreme is defined as having greater absolute log-enrichment). *Enrichment regions.* Seven diverse region definitions were used to assess mutation enrichment. (1) The ChromHMM 15-state model from dorsolateral prefrontal cortex (ID: E073), which assigns every genomic location one of 15 states^46^; (2) transcribed and untranscribed genomic regions as determined by the GENCODEv26 gene model (UTRs, exons and introns were defined as transcribed); (3) promoters and enhancers active in neurons^48^; (4) ChIP-seq measurements of histone modifications H3K27ac, H3K4me1, H3K4me3, H3K9ac, H3K9me3 from the Roadmap Epigenomics Project^46^; (5) RNA-sequencing gene expression levels in the prefrontal cortex (BA9) from the GTEx Portal, v8 release^78^; (6) single nucleus RNA-sequencing gene expression levels from excitatory neurons;^35^ and (7) single nucleus ATAC-sequencing of accessible chromatin in excitatory neurons^35^.

#### Single-nucleus RNA sequencing library preparation

For each of the premotor cortex samples of *C9ORF72* ALS brains and prefrontal cortex samples of *C9ORF72* FTD brains, ten thousand neuronal nuclei with TDP-43 pathology (NeuN+, TDP-43-) and without TDP-43 pathology (NeuN+, TDP-43+) were sorted into individual wells of 96-well plates. For each of the premotor cortex samples and prefrontal cortex samples of age-matched normal brains, only ten thousand neuronal nuclei without TDP-43 pathology (NeuN+, TDP-43+) were sorted. Sorted nuclei were then used for droplet generation and sequencing library preparation using the 10X Genomics Next GEM Single Cell 3′ GEM Kit v3.1 and Chromium Controller, following the manufacturer’s manual.

#### Cryptic exon junction analysis

Annotated TDP-43 dependent cryptic junctions were obtained from the previous study and converted from hg38 to hg19 genome assembly^28^. Cryptic exon junction analysis was performed following the recent study with some modifications^29^. BAM files generated by cellranger were filtered to exclusively include reads meeting the criteria set in the snRNA-seq analysis (described above). For neuron type-specific junction analysis, BAM files were split into distinct neuron type-specific BAM files based on each cell’s 10X barcode and corresponding cluster annotation. These filtered BAM files were subsequently analyzed using regtools’ junctions extract function with theparameters “-a 6 -m 30 -M 500000 -s RF” to generate junction files^79^. Junction files were processed using Leafcutter’s leafcutter_cluster_regtools.py with the parameters “-m 10 -p 0.0001” to generate a summary matrix of junction counts across the dataset^80^.Given the limited number of reads supporting individual splice junctions, read counts from two replicates were summed for downstream analysis. Only genes with at least 3 reads supporting the splicing events were included (defined as the sum of constitutive splicing reads and cryptic exonization reads; corresponding to the denominator in the regtools output).

#### TOP1cc flow cytometry assay

Nuclei were isolated from diseased and control brain tissues as described above. Purified nuclei were fixed using 2% paraformaldehyde (PFA) at 4 °C for 15 min, followed by a wash with ice-cold cell buffer (PBS supplemented with 3mM MgCl2). The nuclei were then stained with Alexa Fluor 488-conjugated anti-NeuN antibody (Millipore MAB377X) and Alexa Fluor 647-conjugated anti-TOP1cc antibody (Millipore MABE1084) in cell buffer containing 0.1% Triton-X100 and 1% bovine serum albumin (BSA) at 4 °C for 1 hr. DAPI was added to stained nuclei immediately before flow cytometry analysis. NeuN+/TOP1cc+ nuclei were analyzed using a BD FACSAria II.

#### TOP1cc RADAR assay

Nuclei from diseased and control brain tissues were isolated as described above, with Halt™ Protease Inhibitor Cocktail (Thermo Fisher 87785) included in all buffers. For each case, 5 x 10^5^ nuclei were sorted into 300 μL of ice-cold PBS supplemented with 3 mM MgCl in 1.5-mL DNA LoBind® tubes (Eppendorf 022431021). Sorted nuclei were centrifuged at 900x g for 10 min at 4 °C, and the supernatant was carefully removed without disturbing the nuclear pellet. The pellet was lysed by adding 1 mL of DNAzol (Thermo Fisher 10503027) containing 1% N-lauroylsarcosine sodium salt (MilliporeSigma L7414), freshly prepared at room temperature. The nuclei were mixed by pipetting 10 times and maintained on ice for all subsequent steps. Glycogen (20 μg; Thermo Fisher R0561) was added to each sample, followed by thorough mixing. Each sample was then equally divided into two 1.5-mL DNA LoBind® tubes. DNA was precipitated by adding 1 mL of ice-cold 100% ethanol and incubating at -20 °C for 10 min, followed by centrifugation at maximum speed for 10 min at 4 °C. The supernatant was carefully removed, and the DNA pellet was washed twice with ice-cold 80% ethanol, with centrifugation at maximum speed for 10 min at 4 °C after each wash. Pellets were air-dried for 3 min at room temperature and resuspended in 100 μL of 8 mM NaOH. Immediately after resuspension, 1 μL of 1 M HEPES buffer was added to neutralize the solution. DNA concentrations were measured using the Qubit™ 1X dsDNA High Sensitivity Assay Kit (Thermo Fisher Q33231). For slot blot analysis, 300 ng of DNA from each sample was applied to a nitrocellulose membrane (Bio-Rad 1620115) using the Bio-Dot® SF Microfiltration Apparatus (Bio-Rad 1706542) according to the manufacturer’s instructions. Membranes were UV-crosslinked for 10 min and blocked in Intercept® (TBS) Blocking Buffer (LI-COR 927-60001) for 1 h at room temperature with gentle rocking. Membranes were incubated with anti-TOP1cc antibody (Millipore MABE1084) or anti-ds DNA antibody (Abcam ab27156) diluted in Intercept® (TBS) Blocking Buffer for 2 h at room temperature. After three washes in TBST (10 min each), membranes were incubated with LI-COR IRDye® secondary antibodies and imaged using the LI-COR Odyssey Imaging System (model 9120). Signal intensities were quantified using ImageJ.

#### META-CS library preparation

META-CS libraries were made following the previously described protocol with modifications (DOI: http://dx.doi.org/10.17504/protocols.io.6qpvr3nbzvmk/v1)^49^. Fifty neuronal nuclei were sorted into individual wells of 96-well plates containing 2 μL of META lysis buffer (20 mM Tris, pH 8.0, 20 mM NaCl, 0.15% Triton X-100, 25 mM dithiothreitol, 1 mM EDTA, 3 units/mL NEB Thermolabile Proteinase K [P8111S]) at 30 °C for 1 h, 55 °C for 10 min. Lysed cells were stored at -80 °C until library preparation.

The transposome was assembled using Tn5 transposase (Diagenode C01070010) following manufacturer’s manual (5 μL of annealed META mix, 5 μL of Tn5, 5 μL of glycerol). The assembled transposome was further diluted using the Tagmentase Dilution Buffer (Diagenode C01070011) based on optimized Tn5 concentration (performed on the day of library preparation). After thawing on ice, cell lysate was transposed with 8 μL of transposition mix (1 μL of diluted transposome, 5 μL of 2X Tagmentation Buffer [Diagenode C01019043], 2 μL of water) and incubated at 55 °C for 15 min. The transposition was stopped by adding 2 μL of 6X stop buffer (300 nM NaCl, 45 mM EDTA, 0.01% Triton X-100, 3 units/mL Thermolabile Proteinase K [NEB P8111S]) and incubated at 37 °C for 30 min, 55 °C for 10 min.

First-strand tagging was performed by adding 13 μL of Strand Tagging Mix 1 (5 μL of Q5 reaction buffer, 5 μL of Q5 high GC enhancer, 0.85 μL of 100 μM [total] Adp1 primer mix, 0.6 μL of 100 mM MgCl2, 0.6 μL of water, 0.5 μL of 10 mM dNTP mix, 0.25 μL of 20 mg/mL bovine serum albumin [NEB B9200], 0.25 μL of Q5 DNA polymerase [NEB M0491]) and incubated at 72 °C for 3 min, 98 °C for 30 s, 62 °C for 5 min, 72 °C for 1 min. Adp1 primers were removed by adding 1 μL of Thermolabile ExoI (NEB M058) and incubated at 37 °C for 15 min, 65 °C for 5 min.

Second-strand tagging was performed by adding of 4 μL of Strand Tagging Mix 2 (1 μL of Q5 reaction buffer, 1 μL of Q5 high GC enhancer, 0.95 μL of 100 μM [total] Adp2 primer mix, 0.945 μL of water, 0.1 μL of 10 mM dNTP mix, 0.05 μL of Q5 DNA polymerase) and incubated at 98 °C for 30 s, 62 °C for 5 min, 72 °C for 1 min. Adp2 primers were removed by adding 1 μL of Thermolabile ExoI (NEB M058) and incubated at 37 °C for 15 min, 65 °C for 5 min.

Strand tagging products were amplified by adding 19 μL of PCR mix (5 μL of NEBNext Multiplex Oligos Universal Primer, 5 μL of NEB Index Primers [NEB E7335S, E7500S, E7710S, E7730S], 4 μL of Q5 reaction buffer, 4 μL of Q5 high GC enhancer, 0.4 μL of 10 mM dNTP mix, 0.4 μL of water, 0.2 μL of Q5 DNA polymerase) and incubated at 98 °C for 20 s, 11 cycles of (98 °C for 10 s, 72 °C for 2 min), 72 °C for 2 min.

Size selection of META-CS libraries were performed using AMPure XP beads (Beckman Coulter A63882) with a 0.65X upper cut and a 1.8X lower cut following manufacturer’s manual. Libraries were eluted into 30 μL of low TE buffer. Each META-CS library was sequenced on one lane of the Illumina HiSeq X sequencer.

#### Strand break assay

gDNA was extracted from disease-relevant brain regions and cerebellum of *C9ORF72* ALS, *C9ORF72* FTD, AD, and control brains using the Qiagen EZ1 Advanced XL system (Qiagen 9001874) in a single batch. For each sample, two parallel reactions were prepared: an S1 nuclease–treated reaction and a matched buffer-only control. In the S1-treated reaction, 400 ng of gDNA was digested with 10 U of S1 nuclease (Thermo Fisher, EN0321) in the presence of 6 μL of 5× reaction buffer in a final volume of 30 μL. Control reactions contained the same components but without S1 nuclease. Reactions were incubated at 37°C for 30 min and subsequently terminated by adding 2 μL of 0.5 M EDTA followed by incubation at 70°C for 10 min. Following digestion, 1 μL of each reaction was analyzed using an Agilent TapeStation with the Genomic DNA ScreenTape assay to quantify DNA fragmentation based on smear intensity. In parallel, 30 μL of each reaction was separated on an agarose gel and visualized by SYBR™ Gold (Thermo Fisher, S11494) staining for 30 min. Smear intensity observed in the buffer-only control reflects pre-existing DSBs, whereas the increase in smear intensity in S1-treated samples relative to controls indicates the presence of SSBs, which are converted into SSBs by S1 nuclease digestion.

#### Preprocessing of META-CS data

We preprocessed the META-CS data using a pipeline that expands the previously reported workflow^49^ with additional features to accommodate both our modified experimental protocol and the somatic Indel calling method. First, paired-end reads were preprocessed by customized pre-meta which identifies Tn5 barcodes, merges overlapping read ends, and trims Illumina adapters. Then, BWA-MEM (v.0.7.17)^81^ was used to map reads to the human reference genome (GRCh37 with decoy) and generate a BAM file for each sample. Importantly, we then split the BAM file by unique pairs of barcodes which are used to distinguish original DNA fragments and help ensure that mutation calling was performed on a single-molecule level. Furthermore, given a relatively small pool of unique barcodes, barcode collision may occur where different DNA fragments are tagged by the same barcode pair around the same genomic region. This possibility also increases with our pooled-cell input with many more available DNA molecules at any locus. To address this challenge, we extracted Tn5 cut sites of each molecule (i.e. the start and end positions of each read pair) and used a combination of shared Tn5 barcode pair and Tn5 cut sites to identify original DNA fragments.

#### Somatic Indel calling from META-CS data

As the original META-CS method was developed for somatic SNVs, we established a somatic Indel calling pipeline (https://github.com/gldong/duplex-indel) that introduces modules to enhance the calling accuracy and remove false positives specific for Indels^82^. First, we followed a workflow similar to SNV calling and piled up reads from the META-CS-barcode pair BAMs and a matching bulk WGS BAM to generate somatic Indel candidates by identifying variants that have at least 4 total non-reference (ALT) reads in META-CS with at least 2 ALT reads from each strand for duplex support, as well as no ALT read in bulk to remove potential germline variants.

When mate-pair reads from the same DNA molecule overlapped, the two mates were merged into a single read to avoid double counting. However, merging across repetitive DNA regions can introduce alignment artifacts and create spurious Indel evidence. To reduce these false positive calls, we defined a merging window as the overlapping region between mate-pair reads, and any Indel intersecting the merging window was discarded. When reads had different overlap boundaries, the largest merging window was used by taking the smallest start position and the largest end position.

In addition, we implemented a set of filters. A candidate site was filtered out if it met any of the following criteria, 1) overlapping with the low-quality regions as previously reported (um75-hs37d5.bed^49^), 2) overlapping with gnomAD Indels of ≥ 1% population frequency, 3) located within 100 bp from another candidate site, 4) located within 10 bp of read ends, 5) having multiple Tn5 cut sites.

#### Single-strand Indel calling from META-CS data

Similar to double-strand Indel calling described above, single-strand Indel candidates were generated by piling up reads from the META-CS barcode-pair BAMs and a matching bulk WGS BAM, followed by identifying variants that have no ALT read from bulk and at least 8 total reads in META-CS with at least 4 reference allele (REF) reads from one strand (i.e. non-variant strand) and at least 4 ALT reads from the other strand (i.e. variant strand). In addition, candidates are required to have no ALT reads from the non-variant strand and no REF reads from the variant strand. Other filters are the same as double-strand calling above.

#### ALS and FTD snRNA-seq analysis

For snRNA-seq analysis on ALS and FTD, we used data generated for this study (i.e. in-house). We implemented the following pipeline to raw FASTQ files from both data sets. Cellranger (v.6.1.0)^83^ with “--include-introns” was used to map reads to the human reference genome (GRCh37) and generate gene-by-cell count matrices. To remove potential ambient RNA contamination, we ran remove-background from CellBender (v.0.3.0)^84^on the raw count matrices with the default parameters to generate filtered count matrices for downstream analysis. For quality control, we filtered out cells with < 700 genes and mitochondrial gene percentage > 20%, which resulted in more than 65% cells removed in one sample (1700BA6-normal). So this sample was excluded for further analysis. scDblFinder^85^was used to remove doublets, and DropletQC^86^was used to calculate nuclear fractions where cells with a nuclear fraction < 0.25 were filtered out. Next, we used Seurat (v.5.1.0)^87^ SCTransform v2 to perform normalization and variance stabilization followed by RunHarmony^88^ to integrate and harmonize cell embeddings across all samples. The harmonized embeddings were used to perform dimension reduction (RunUMAP) and clustering using the Leiden algorithm (resolution = 0.8)^89^. Cell types were manually annotated using previously reported markers^90^ and canonical markers. Specifically, each marker gene’s expression was visualized on the UMAP, and clusters with the highest and most specific expression were assigned to the corresponding cell type. A general cell type (excitatory neurons, inhibitory neurons, and non-neuronal cells) was annotated first, followed by subtyping of neurons. For example, clusters exhibiting high expression of SLC17A7 were annotated as excitatory neurons, and subclusters demonstrating concurrent high expression of CUX2 were further annotated as L2-3 IT.

#### Cell type fraction analysis in snRNA-seq

We used a Bayesian statistics framework^91,92^ to detect changes in cell type fractions in ALS and FTD compared to controls. This model implements Dirichlet multinomial modeling – Hamiltonian Monte Carlo (DMM-HMC) using R packages rstan (v.2.32.6) and bayestestR (v0.13.2) (http://mc-stan.org/). The input was a count matrix where each row is a sample and each column is a cell type. Counts in each row of the matrix follow a multinomial distribution where proportions of each cell type in the sample are modeled by Dirichlet. The prior probability of the proportions is another Dirichlet distribution with the fixed parameter alpha = 10^-7^ for equal expected fractions of cell types. We split samples into groups based on their disease status, brain region, and TDP-43 status, and calculated the posterior probability distribution (PPD) of each group. Then, we subtracted the PPD of control group from that of disease group to detect any significant changes in cell type fractions. Significance was determined if the log2 ratio of disease/control has 95% credible intervals above or below zero.

#### Differential gene expression analysis in snRNA-seq data

Differential gene expression analysis was performed using Nebula (v.1.5.3)^93^ in R on in-house ALS and FTD data set and AD data set from SEA-AD consortium^94^. We downloaded the processed AD snRNA-seq data from SEA-AD consortium’s web portal (sea-ad.org). For each data set, raw count matrices after preprocessing and cell type annotation were used as inputs and the design matrix was constructed with disease status as the predicator. Then, a negative binomial mixed model was built for each gene using individual as the random effect and total number of molecules in each cell as offsets. The output raw p-values were adjusted using the Benjamini Hochberg method. Differentially expressed genes with FDR < 0.05 were considered significant.

#### qPCR analysis of RNASEH2 and RRM gene expression

Neuronal nuclei (2350 NeuN+ nuclei, ∼5 μL) from *C9ORF72* FTD, AD and age-matched control brains were sorted into individual wells of 96-well plates with triplicates. Cell lysis, reverse transcription and qPCR were conducted using the Power SYBR™ Green Cells-to-CT™ Kit (Thermo Fisher 4402954) following the manufacturer’s protocol. The following primers were used for qPCR analysis:

hRNASEH2A Forward - 5’- ACAGCCACTGGGCTTATACAG -3’

hRNASEH2A Reverse - 5’- TCCCTACGGTGTCCACGAATA -3’

hRNASEH2B Forward - 5’- TAACCCCTGTTCAGGAGAAGG -3’

hRNASEH2B Reverse - 5’- ACACGTTATCCACCACAACTTG -3’

hRNASEH2C Forward - 5’- AGGGACTCGAAGTGTCGTTTC -3’

hRNASEH2C Reverse - 5’- TGTCACCATCACGTATCCCAC -3’

hRRM1 Forward - 5’- GCCAGGATCGCTGTCTCTAAC -3’

hRRM1 Reverse - 5’- GAGAGTGTTTGCCATTATGTGGA -3’

hRRM2 Forward - 5’- ATTGGGCCTTGCGATGGATAG -3’

hRRM2 Reverse - 5’- GAGTCCTGGCATAAGACCTCT -3’

hRBFOX3 Forward - 5’- TCGTAGAGGGACGGAAAATTGA -3’

hRBFOX3 Reverse - 5’- GCCGTTGGTGTAGGGGTTC -3’

#### TDP-43 knockdown in NGN2 neurons

CRISPR inhibition (CRISPRi) mediated knockdown of TDP-43 was performed in KOLF2.1J and FA11 iPSC lines harboring Tetracycline-inducible Neurogenin2 (NGN2) transcription factor and dCAS9-ZIM3-KRAB-BFP. NGN2-mediated differentiation of iPSCs was achieved as previously described^95^. Briefly, undifferentiated iPSCs were maintained in StemFlex media (Gibco) with 1X StemFlex supplement (Gibco). On Day 0 of differentiation, cells were plated at a density of 50,000 cells/cm2 in induction media (97% DMEM/F12 + 1% N2 supplement, 1% Non-Essential Amino Acids, 1% Glutamax, 2 ug/mL doxycycline), and transduced with lentivirus expressing either non-targeting dual guide RNAs or TARDBP-targeting dual guide RNAs. Successfully transduced cells were selected with 5ug/ml puromycin for 48 hours. On Day 3 of differentiation, cells were re-plated on 0.1% poly-L-ornithine- and 5ug/ml laminin-coated-plates in differentiation media (95% Neurobasal medium + 2% B27 supplement, 1% Glutamax, 50mM sodium chloride, 10 ng/ml NT3, 10 ng/mL BDNF, 10 ng/mL GDNF, 5uM Aphidicolin, 2 ug/mL doxycycline). Half-media changes were done every 3 days and cells were harvested on Day 10 of differentiation for either DNA extraction or immunofluorescence.

#### Immunofluorescence protocol

iPSC-derved neurons cultured for 10 days in 96-well plates were fixed in 4% parafolrmadehyde for 20 minutes at room temperature. Neurons were permeabilized in 0.2% Triton X-100 diluted in PBS for 10 minutes followed by blocking in 5% bovine serum albumin diluted in PBS for 1 hour at room temperature. Neurons were incubated in primary antibodies diluted in 5% BSA for 2 hours at room temperature, followed by secondary antibodies diluted in 5% BSA for 1 hour at room temperature. Neurons were counterstained with DAPI and imaged at 40x magnification on Nikon C1 confocal microscope. Primary antibodies: anti-TDP-43 (rabbit polyclonal, Proteintech 10782-2-AP, 1/500 concentration) and anti-MAP2 (chicken polyclonal, Thermo Fisher PA1-10005, 1/5000 concentration). Secondary antibodies: Donkey anti-rabbit Alexa Fluor 488 (Thermo Fisher A21206) and Donkey anti-chicken Alexa Fluor 647 (Thermo Fisher A78952).

### QUANTIFICATION AND STATISTICAL ANALYSIS

All of the quantitative and statistical methods, strategies, and analyses are described in the relevant sections of the method details or in the table and figure legends.

---

**Figure S1.**
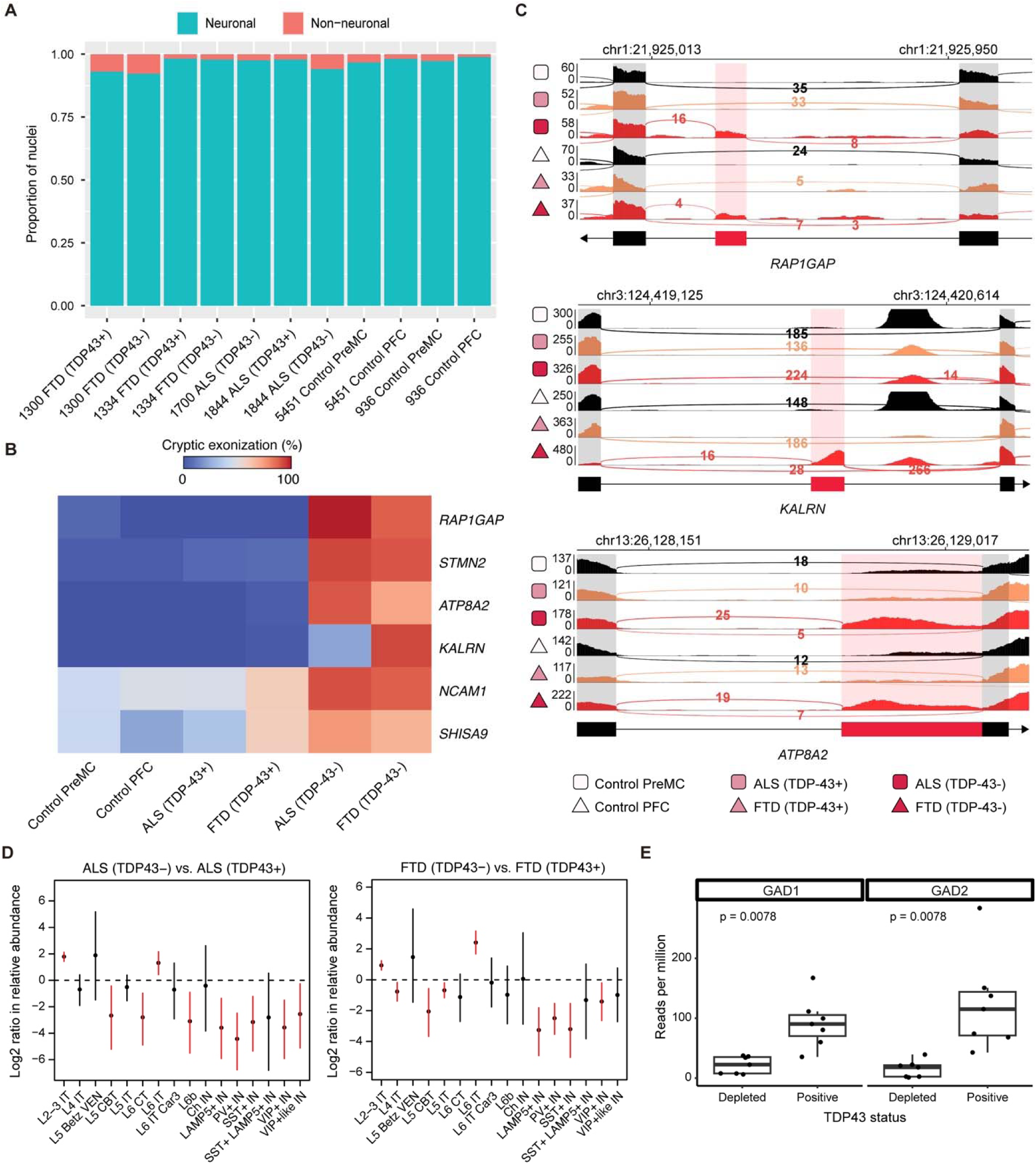
Characterization of isolated TDP-43+ and TDP-43- neuronal nuclei from *C9ORF72* ALS and FTD brains, related to Figure 1. (A) Proportion of neuronal and non-neuronal nuclei across different cell and case conditions. Bars represent the fraction of neuronal (blue) and non-neuronal (red) nuclei for each dataset. (B) Heatmap showing the fraction of transcripts containing cryptic exons across various genes in different cell and case conditions. (C) IGV Genome browser tracks of *RAP1GAP*, *ATP8A2* and *KALRN* with snRNA-seq read coverage and splice junctions. TDP-43- neurons (red) show an increase in cryptic exon inclusion (highlighted region), which is low or absent in TDP-43+ (orange) and control (black) neurons. Numbers adjacent to splice junctions represent read counts supporting the junction. (D) Log fold change in neuronal subtype abundance in TDP-43- vs. TDP-43+ neurons. Error bars show 95% credible intervals of posterior distributions from a Dirichlet multinomial model (see STAR Methods). Red intervals indicate significant changes in neuronal subtype abundance. IT: intratelencephalic neurons. VEN: Von Economo neurons. CBT: corticobulbar tract neurons. CT: corticothalamic neurons. PV: parvalbumin. IN: inhibitory neurons. Ch IN: cholinergic inhibitory neurons. (E) Expression of inhibitory neuron marker genes in TDP-43- neurons. Box plots compare reads per million for *GAD1* and *GAD2* expression between TDP-43- and TDP-43+ neurons from *C9ORF72* FTD brains using previously generated bulk RNA-seq data. *P*-values are one-tailed paired Wilcoxon rank-sum tests.

**Figure S2.**
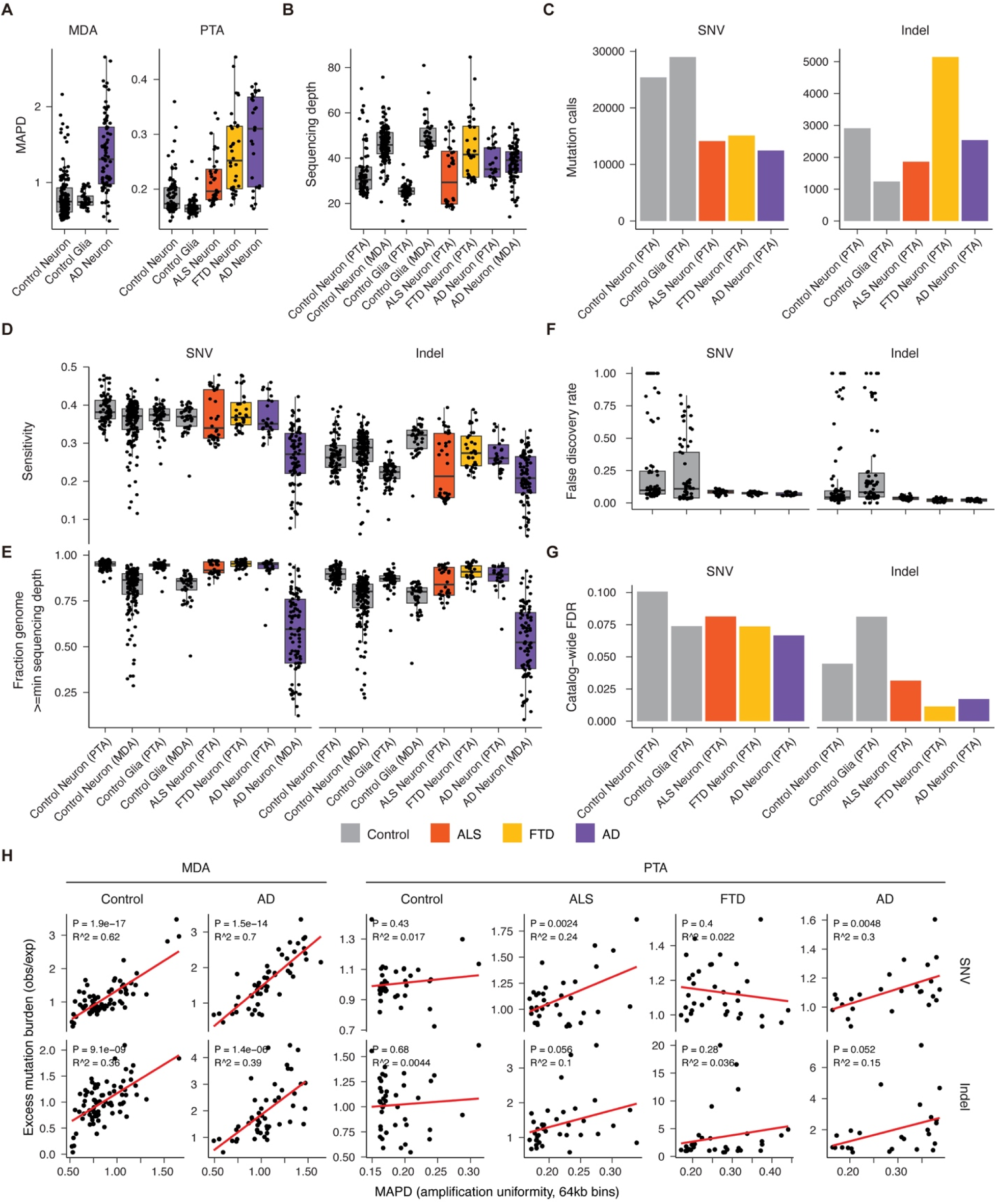
scWGS metrics and quality assessment for PTA-amplified and MDA-amplified single cells, related to Figure 2. (A) Median Absolute Pairwise Difference (MAPD), a measure of amplification uniformity; low values indicate more uniform amplification. Glia include pure oligodendrocytes and mixed glia^35^. (B) Mean sequencing depth per cell. (C) Total numbers of somatic mutations called by SCAN2. (D) SCAN2 mutation detection sensitivity. (E) Fraction of the genome that passes the minimum read depth requirement for SCAN2. Different minimum read depths are used for sSNVs and sIndels. (F) Estimated false discovery rate per single cell based on a previously published false positive rate from simulated data^26^. (G) Estimated false discovery rate for each mutation catalog. That is, the sum of estimated false positives across all cells depicted in (F) divided by the total catalog sizes in (C). In all panels, points represent single cells. (H) Amplification non-uniformity does not explain increased somatic indel burdens in PTA-amplified single neurons. Amplification non-uniformity is measured by MAPD (x-axis), where larger values indicate more uneven amplification. While mutation burden was significantly correlated with MAPD in MDA-amplified cells, somatic indel burdens in PTA-amplified neurons were not significantly associated at the *P* = 0.05 level. Points correspond to individual cells; red lines are simple linear regressions; *P*-values are uncorrected for multiple hypothesis testing.

**Figure S3.**
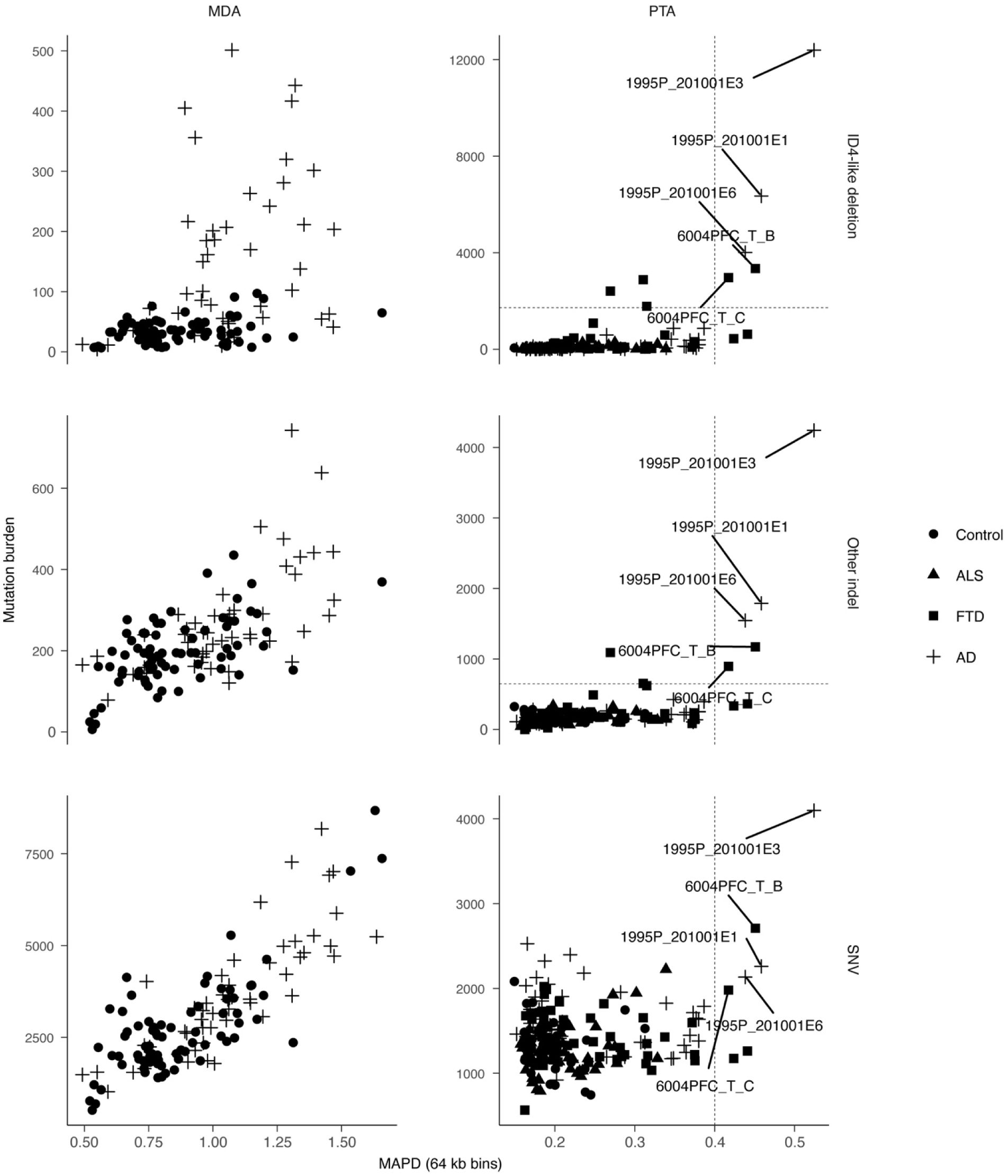
Identification of five PTA-amplified outlier cells, related to Figure 2. Mutation burdens are split into three types: 2-bp ID4-like deletions, all other insertions and deletions, and SNVs. Burdens (which by definition are scaled for mutation detection sensitivity) are plotted against MAPD, a measure of amplification non-uniformity for which higher values indicate less uniform amplification. Outliers were visually identified on the basis of having both high MAPD and high somatic indel burdens. Outlier cutoffs are shown by dashed line. All five outlier PTA cells are labeled. None of these outlier cells were included in any analysis.

**Figure S4.**
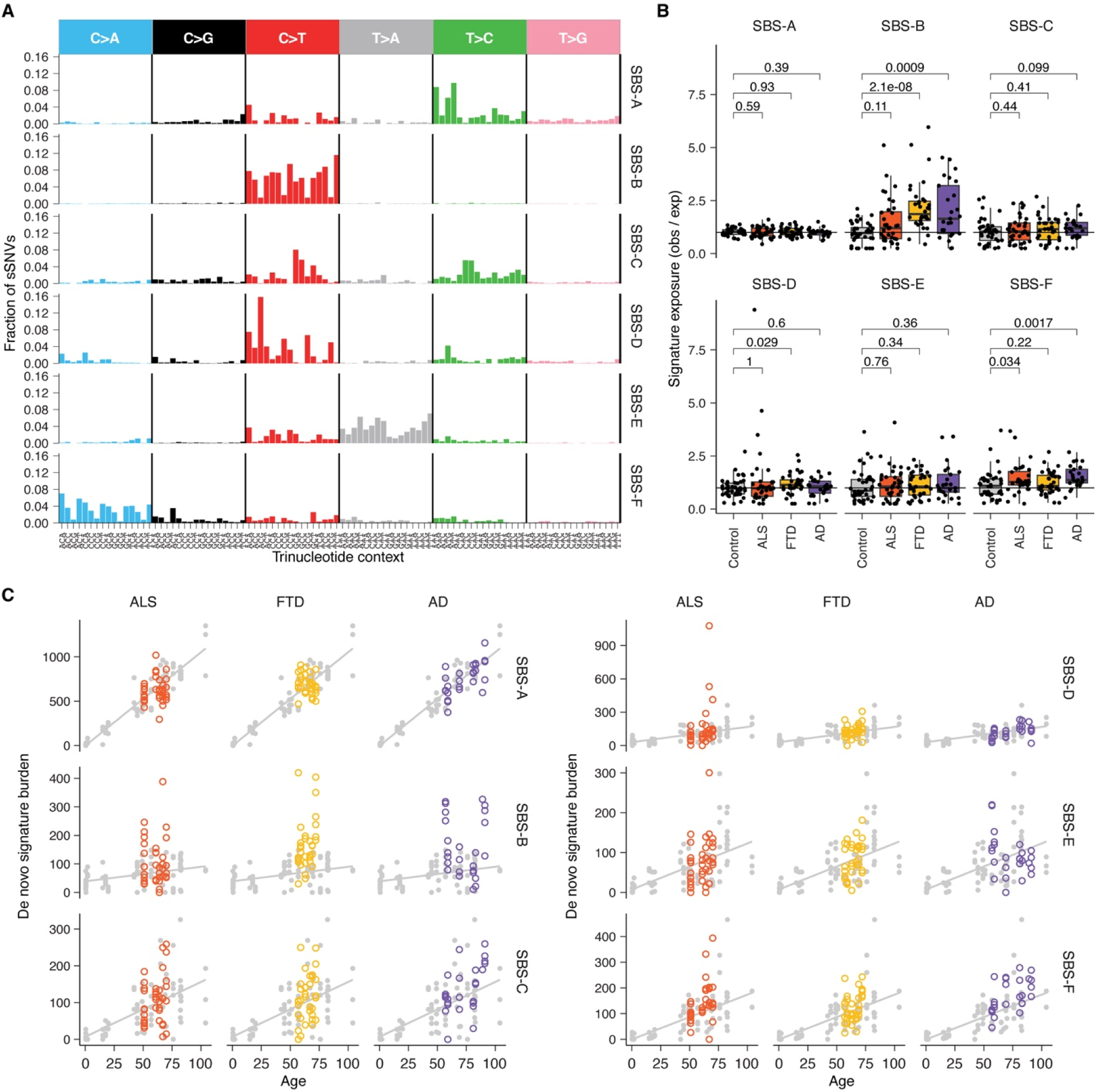
*De novo* SNV signature burdens in PTA-amplified neurons, related to Figure 3. (A) *De novo* SBS signatures identified from joint analysis of diseased and neurotypical control neurons amplified by PTA as well as previously published neurons and oligodendrocytes amplified using an older amplification technology (MDA). MDA data were used only to extract de novo signatures; none of the data points on this figure represent MDA-amplified cells. The x-axis shows the 96 possible trinucleotide mutation contexts, grouped by the six possible base changes, accounting for reverse complements. The y-axis represents the fraction of sSNVs attributed to each signature. (B) Observed SBS signature exposure divided by the age-adjusted expected SBS signature exposure derived from control neurons (see STAR Methods, and panel C). For burden comparison to control neurons, only observed/expected from individuals with age > 50 were included to match the age distribution of disease cases. *P*-values represent Wilcoxon rank-sum tests. (C) Burdens of *de novo* SBS signatures in neurons vs. age. Points and regression lines in gray indicate the age-specific expected SBS exposure derived from control neurons (see STAR Methods) and are identical in each row.

**Figure S5.**
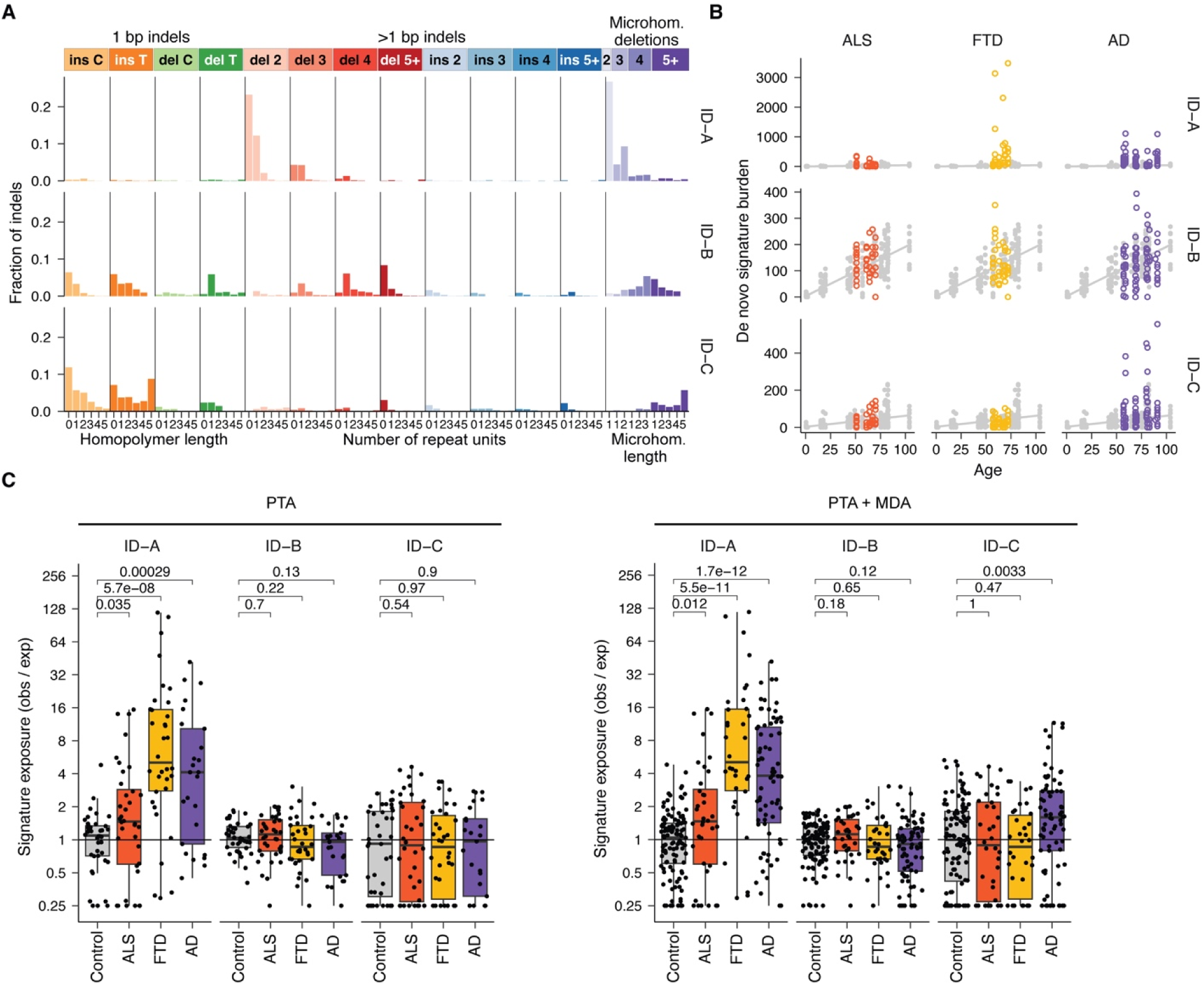
*De novo* sIndel signature analysis in PTA- and MDA-amplified neurons, related to Figure 3. (A) *De novo* sIndel signatures identified from joint analysis of diseased and neurotypical control neurons amplified by PTA as well as previously published neurons and oligodendrocytes amplified using an older amplification technology (MDA). Unlike SNVs, sIndel calling from PTA- and MDA-amplified neurons were generally similar, suggesting that MDA data are useful for quantifying sIndel effects. The x-axis shows the Indel classes defined by insertion/deletion type, Indel length, and local sequence context. The y-axis represents the fraction of sIndels attributed to each signature. (B) Burdens of *de novo* ID signatures in neurons vs. age. Points and regression lines in gray indicate the age-specific expected ID exposure derived from control neurons (see STAR Methods) and are identical in each row. MDA and PTA-amplified neurons are shown. (C) Observed Indel signature exposure divided by the age-adjusted expected Indel signature exposure derived from control neurons (see STAR Methods and panel B). For burden comparison to control neurons, only observed/expected from individuals with age > 50 were included to match the age distribution of disease cases. *P*-values represent Wilcoxon rank-sum tests. Left: PTA-only data. Right: combined PTA and MDA data.

**Figure S6.**
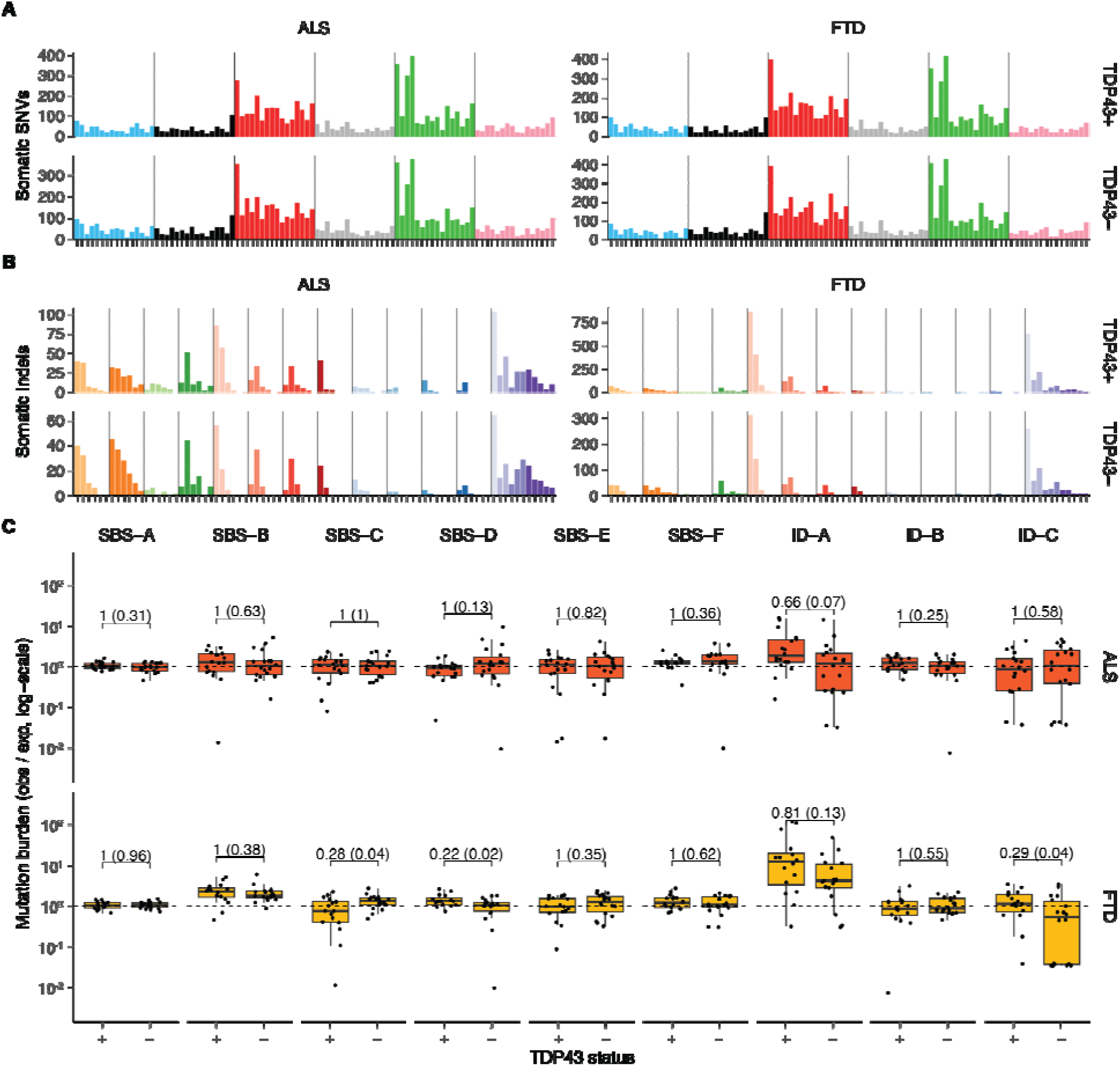
The mutational spectra of TDP43-depleted neurons do not significantly differ from TDP43-positive neurons, related to Figure 3. (A) Summed SNV spectra across all cells. (B) Summed indel spectra across all cells. (C) Excess burden per cell for each of the 9 *de novo* mutational signatures. Values outside of parentheses are Wilcox rank-sum *P*-values corrected for multiple hypothesis (n=9) tests using Holm’s method; values in parentheses are the *P*-values prior to correction.

**Figure S7.**
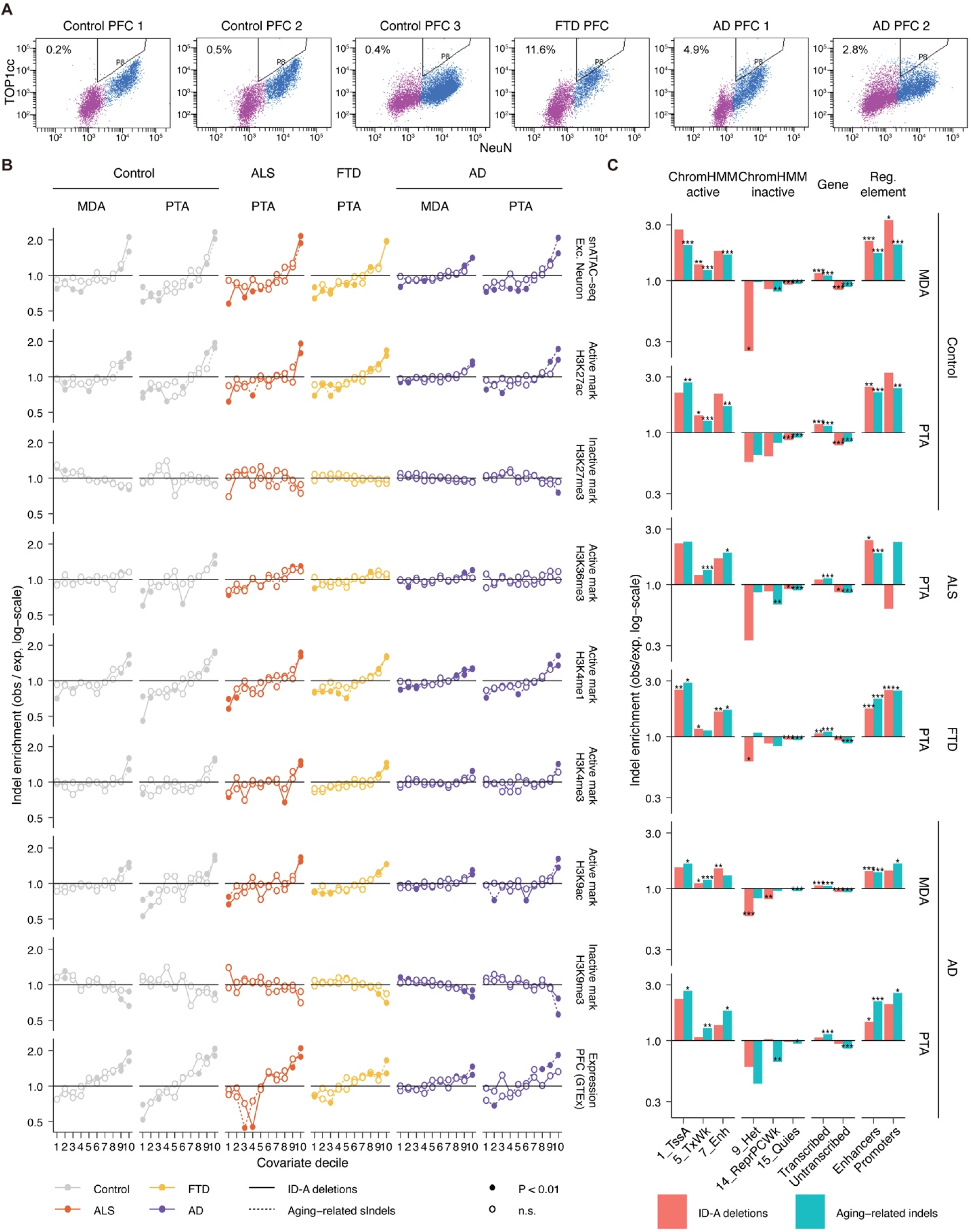
TOP1cc enrichment and genomic features of ID-A-like deletions in neurodegenerative brains, related to Figure 4. (A) Flow cytometry analysis of the TOP1 cleavage complex (TOP1cc) levels in NeuN+ neuronal nuclei from *C9ORF72* FTD, AD and control brains. Gates mark the population of neuronal nuclei with elevated TOP1cc levels, with the percentage of TOP1cc+ neurons indicated in each plot. PFC: prefrontal cortex. (B) sIndel enrichment analyses comparing sIndel burden with GTEx gene expression from prefrontal cortex; local chromatin accessibility in snATAC-seq data from neurotypical excitatory neurons; and histone modifications indicative of active and inactive (H3K9me3) chromatin from prefrontal cortex. Solid lines: ID-A-like sIndels, dotted lines: sIndels associated with normal aging. (C) Enrichment analyses comparing sIndel burdens with chromatin states (determined by ChromHMM^47^) in prefrontal cortex, transcribed and untranscribed genomic regions defined by GENCODE v26, and promoters and enhancer elements active in neurons^48^. Red bars: ID-A-like sIndels, blue bars: sIndels associated with normal aging. Asterisks represent enrichment test P-values (see STAR Methods): * - p < 0.01, ** - p < 0.001, *** p < 0.0001. See Figure 3C for the definition of ID-A-like sIndels.

**Figure S8.**
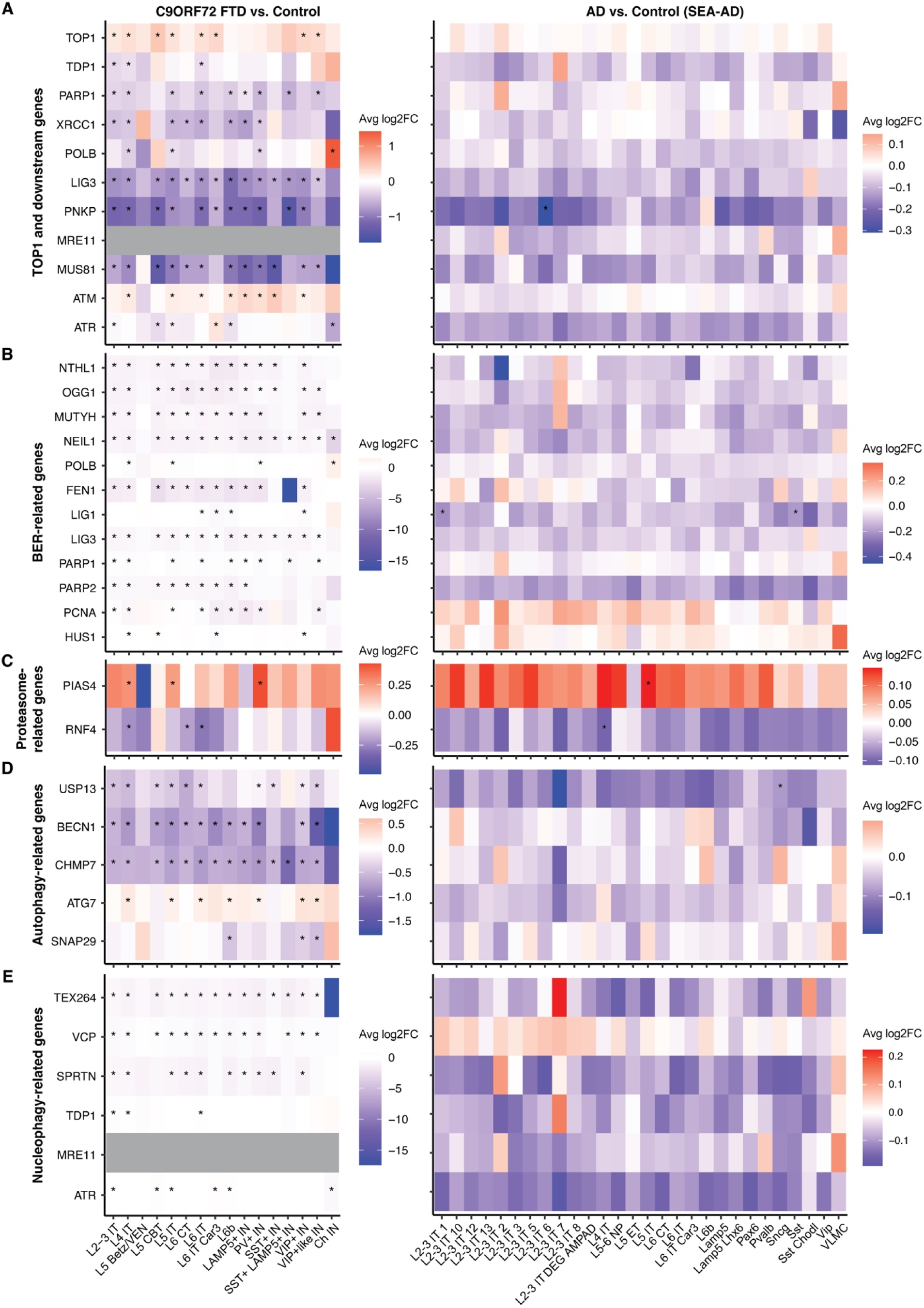
Neuronal expression changes in TOP1 and related DNA repair, proteostasis, autophagy, and nucleophagy pathways in neurodegenerative conditions, related to Figures 4 and 5. (A) Heatmaps showing differential expression of TOP1-related genes in neurons from *C9ORF72* FTD and AD brains compared to controls. (B) Heatmaps showing differential expression of BER-related genes in neurons from *C9ORF72* FTD and AD brains compared to controls. (C) Heatmaps showing differential expression of proteasome-related genes in neurons from *C9ORF72* FTD and AD brains compared to controls. (D) Heatmaps showing differential expression of autophagy-related genes in neurons from *C9ORF72* FTD and AD brains compared to controls. (E) Heatmaps showing differential expression of nucleophagy-related genes in neurons from *C9ORF72* FTD and AD brains compared to controls. The AD vs. control comparisons are based on the Seattle Alzheimer’s Disease Brain Cell Atlas (SEA-AD) dataset. Color scales represent average log FC, with red indicating upregulation and blue indicating downregulation. Grey indicates genes not detected. FDR-adjusted *P*-values < 0.05 are marked by asterisks. IT: intratelencephalic neurons. VEN: Von Economo neurons. CBT: corticobulbar tract neurons. CT: corticothalamic neurons. PV: parvalbumin. IN: inhibitory neurons. Ch IN: cholinergic inhibitory neurons. NP: near-projecting neurons. ET: extratelencephalic neurons. VLMC: vascular leptomeningeal cells.

**Figure S9.**
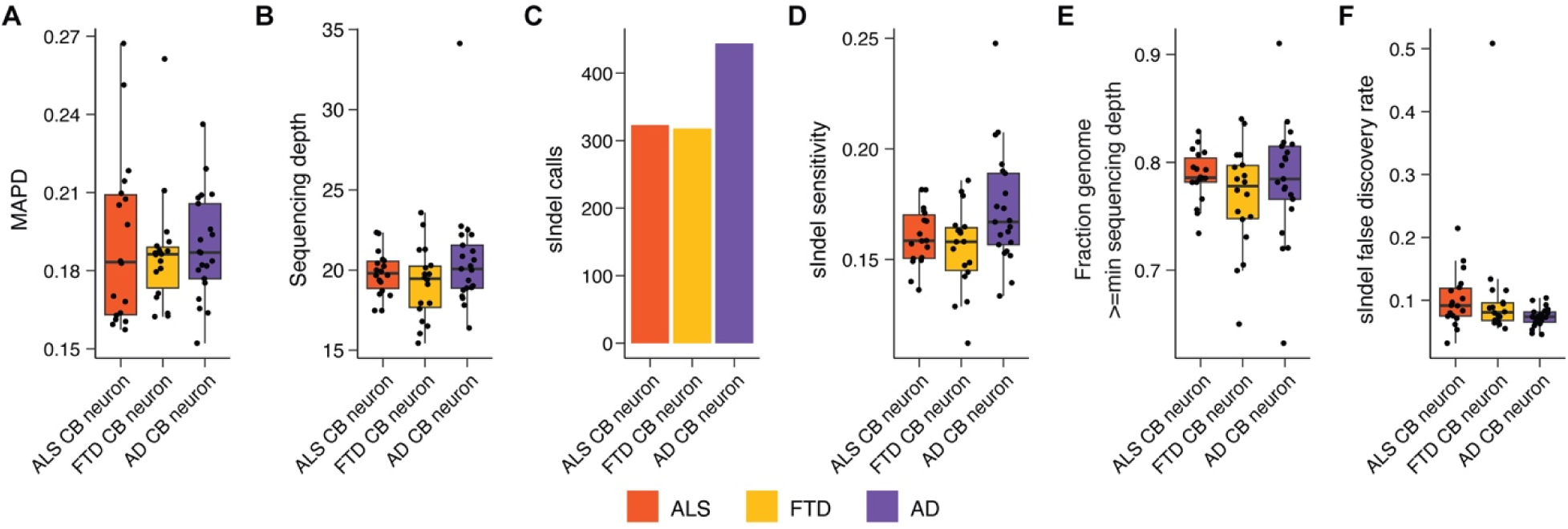
scWGS metrics and quality assessment for PTA-amplified cerebellar neurons, related to Figure 6. (A) Median Absolute Pairwise Difference (MAPD), a measure of amplification uniformity; low values indicate more uniform amplification. CB: cerebellum. (B) Mean sequencing depth per cell. (C) Total numbers of sIndel called by SCAN2. (D) SCAN2 sIndel detection sensitivity. (E) Fraction of the genome that passes the minimum read depth requirement for SCAN2. Different minimum read depths are used for sSNVs and sIndels. (F) Estimated sIndel false discovery rate per single cell based on a previously published false positive rate from simulated data (see STAR Methods)^26^.

**Figure S10.**
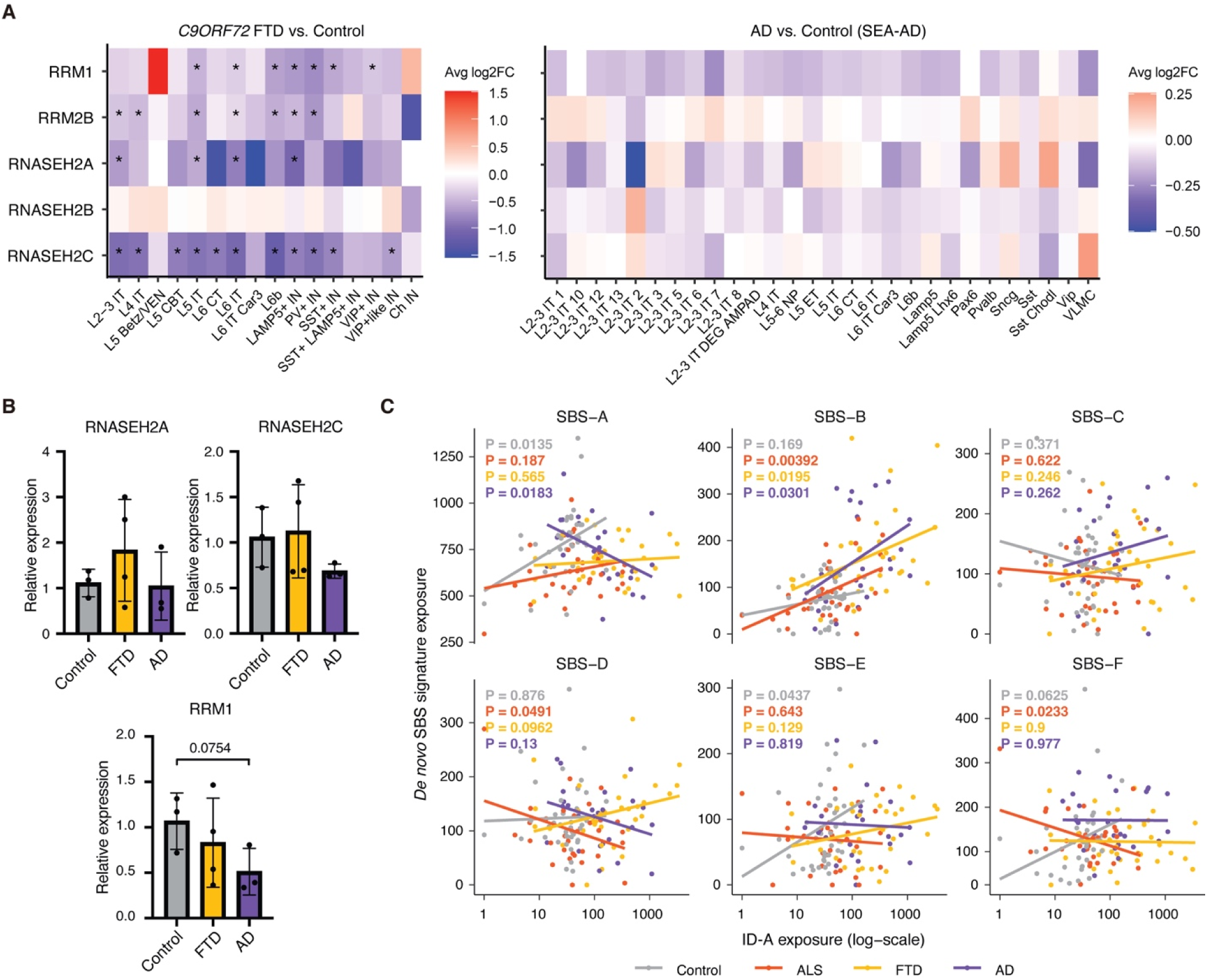
Dysregulation of RNASEH2 and RRM genes and correlation between ID-A and oxidative stress in neurodegenerative conditions, related to Figure 7. (A) Heatmaps showing differential expression of RNASEH2 and RRM genes in neurons from *C9ORF72* FTD and AD brains compared to controls. The AD vs. control comparisons are based on the Seattle Alzheimer’s Disease Brain Cell Atlas (SEA-AD) dataset. Color scales represent average log FC, with red indicating upregulation and blue indicating downregulation. FDR-adjusted *P*-values < 0.05 are marked by asterisks. IT: intratelencephalic neurons. VEN: Von Economo neurons. CBT: corticobulbar tract neurons. CT: corticothalamic neurons. PV: parvalbumin. IN: inhibitory neurons. Ch IN: cholinergic inhibitory neurons. NP: near-projecting neurons. ET: extratelencephalic neurons. VLMC: vascular leptomeningeal cells. (B) Expression levels of *RNASEH2A, RNASEH2C* and *RRM1* in neurons measured by qPCR. *P*-values represent two-tailed unpaired *t*-tests. (C) Correlation between the disease-associated ID-A sIndel signature and all six *de novo* SBS signatures. Each points represents a PTA-amplified cortical neuron. *P*-values are regression *t*-tests on slope, and text color indicates phenotype.

**Figure S11.**
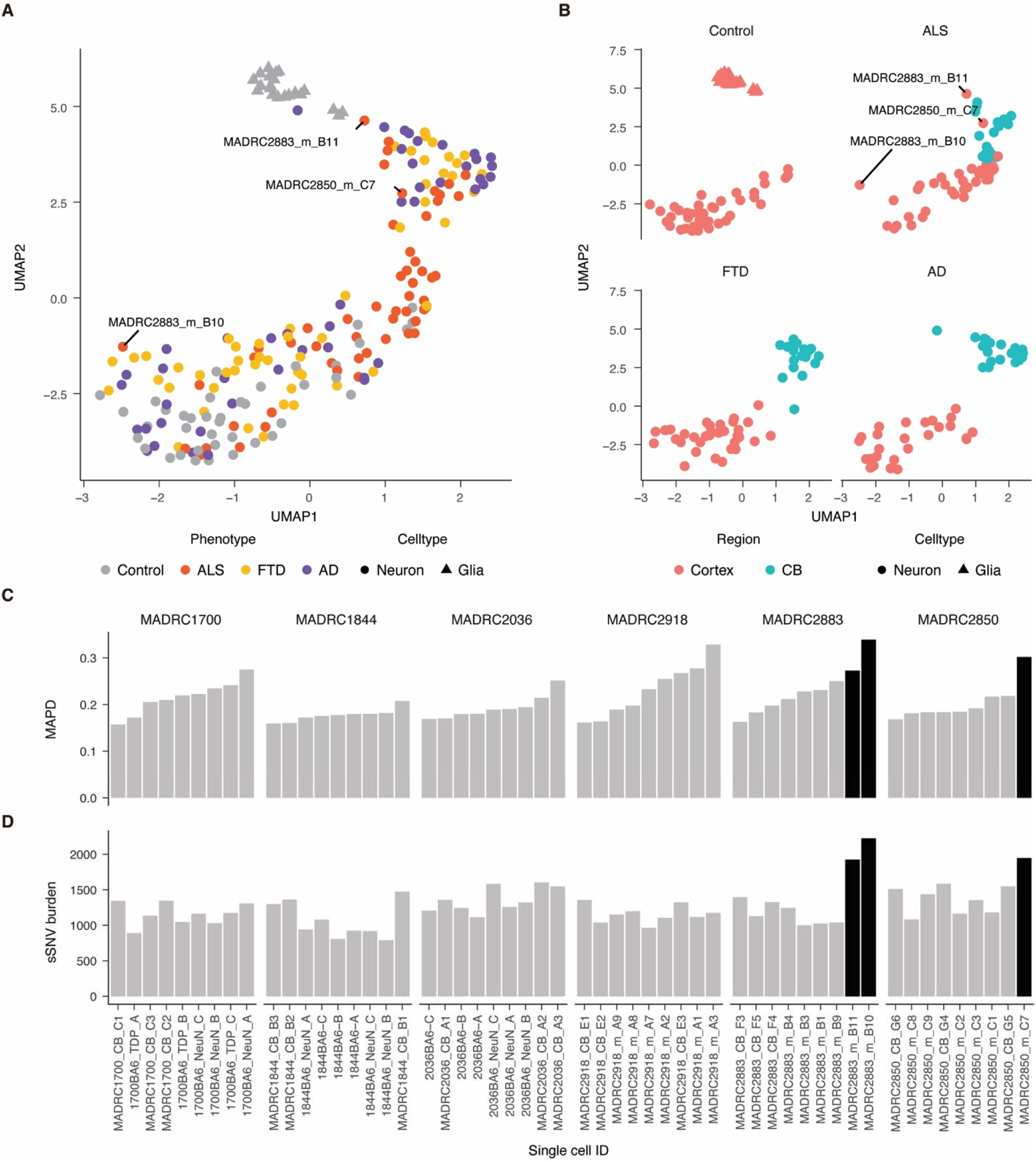
Identification of three outlier PTA-amplified cortical neurons from ALS brains, related to Figure 7. (A) Clustering by UMAP of all PTA-amplified single cells with age > 50, including oligodendrocytes and cerebellar neurons, by their sSNV spectra. Each point represents one cell and is colored by disease phenotype. (B) Same UMAP plots as panel (A) but split by disease phenotype and colored to differentiate cortical neurons (blue) from cerebellar neurons (red). Two outlier cortical ALS neurons cluster with cerebellar ALS neurons, which is not observed in FTD or AD. A third outlier cortical ALS neuron does not cluster with other ALS neurons. (C) Measurements of amplification uniformity by MAPD for all ALS neurons, grouped by donor. Higher MAPD values indicate worse uniformity. The three labeled ALS cells from panels (A) and (B) are shown in black and are elevated compared to other neurons from the same donor. (D) sSNV burdens for all ALS neurons, again grouped by donor and with the outlier cells shown in black.

**Figure S12.**
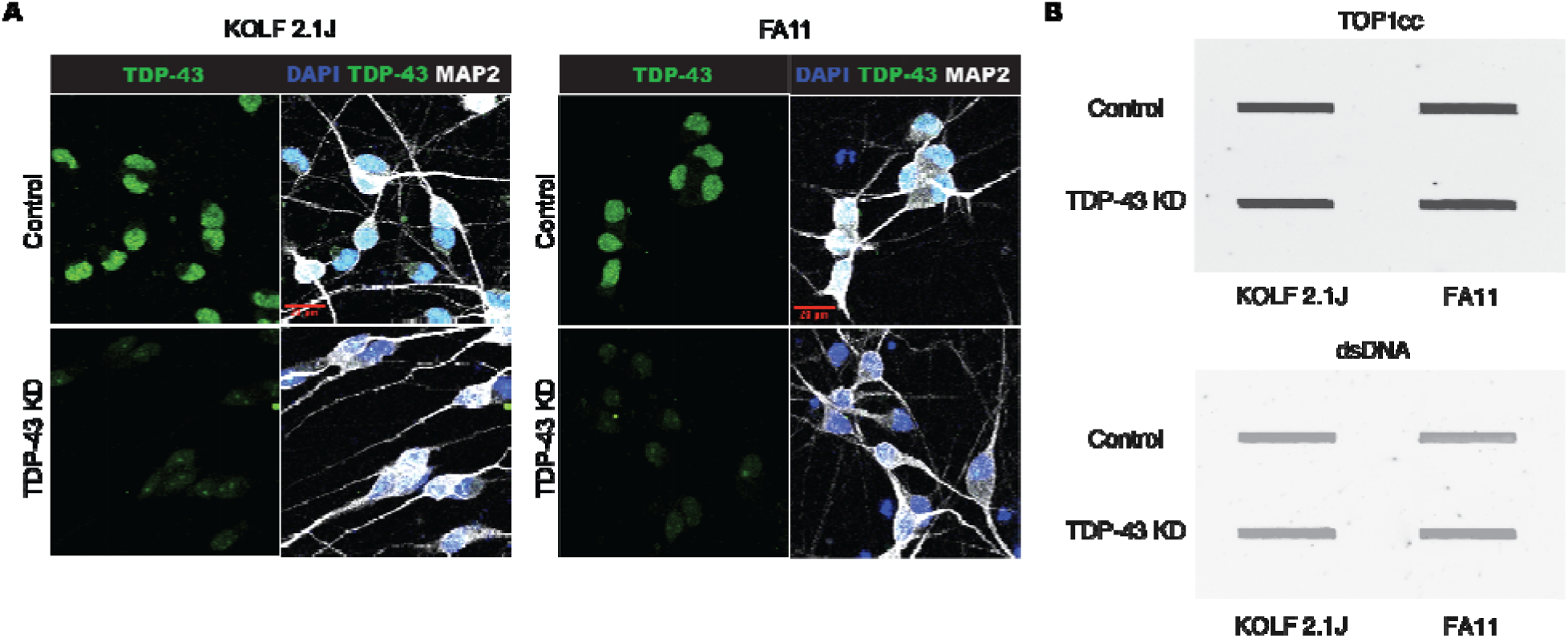
TDP-43 depletion does not induce TOP1cc formation in neurons, related to STAR Methods. (A) Representative images of TDP-43 immunofluorescence (green) in either KOLF2.1J (left) or FA11 (right) iPSC-derived NGN2 neurons. Neurons expressing non-targeting dual guide RNAs show TDP-43 expression in the nucleus (top), while neurons expressing TARDBP-targeting dual guide RNAs show depleted TDP-43 in the nucleus (bottom). Merged panels show MAP2 for differentiated neurons and DAPI for nuclei. Scale bar: 20 _μ_m. (B) Slot blot analysis of TOP1cc (top) and dsDNA (bottom; loading control) in the two NGN2 neuronal cell lines (KOLF 2.1J and FA11). No differences in TOP1cc levels were observed between control and TDP-43 knockdown conditions in either cell line.

**Figure S13.**
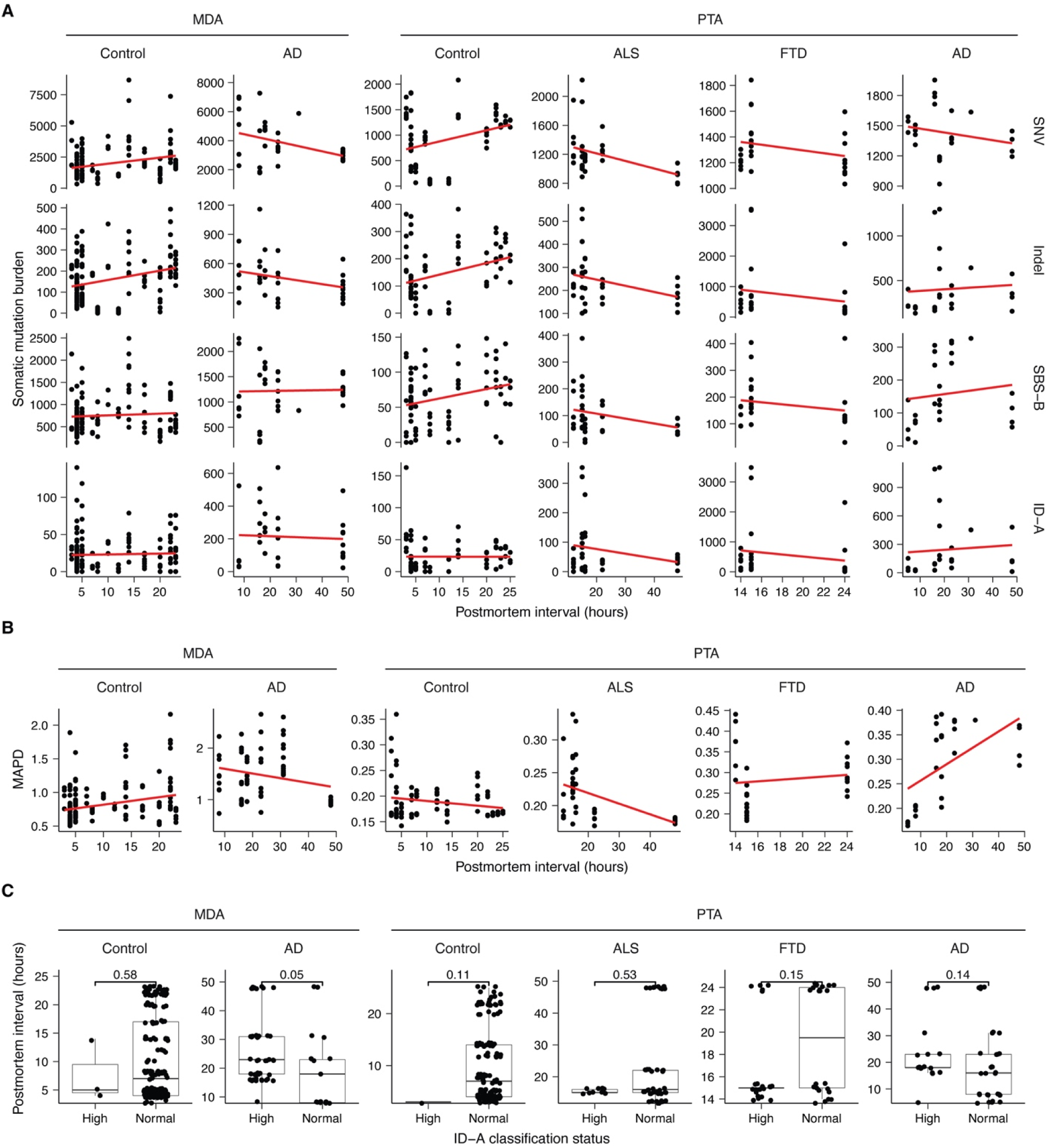
Postmortem interval (PMI) does not drive disease-associated mutation burdens or ID-A-like deletion status in neurons, related to STAR Methods. (A) Postmortem interval (in hours after death) is plotted against somatic mutation burdens of all SNVs, all indels, SBS-B-like SNVs, and ID-A like sIndels. Red lines are simple linear regressions. (B) Postmortem interval (in hours after death) is plotted against a metric of single-cell amplification uniformity, MAPD. Higher MAPD values indicate worse uniformity and thus worse quality. Red lines are simple linear regressions. (C) PMI distributions for cells classified as having typical (ID-A normal) or unusually high (ID-A high) levels of ID-A-like deletions. *P*-values are Wilcoxon rank-sum two-tailed tests. In all plots, each point corresponds to one single cell. Only cortical neurons are shown.

## SUPPLEMENTAL INFORMATION

Table S1. Information of the control and neurodegenerative disease cases and samples, related to Figures 1 and 6

Table S2. Number of cells passing and failing QC across cell types and conditions, related to Figures 2 and 6

Table S3. Metadata of single cells across cell types and conditions, related to Figures 1–3 and 6

Table S4. sSNVs and sIndels identified in single cells listed in Table S3, related to Figures 2–7

Table S5. Burden of sSNVs, sIndels, and mutational signatures in single cells listed in Table S3, related to Figures 2–3 and 6

---

**Data S1.**
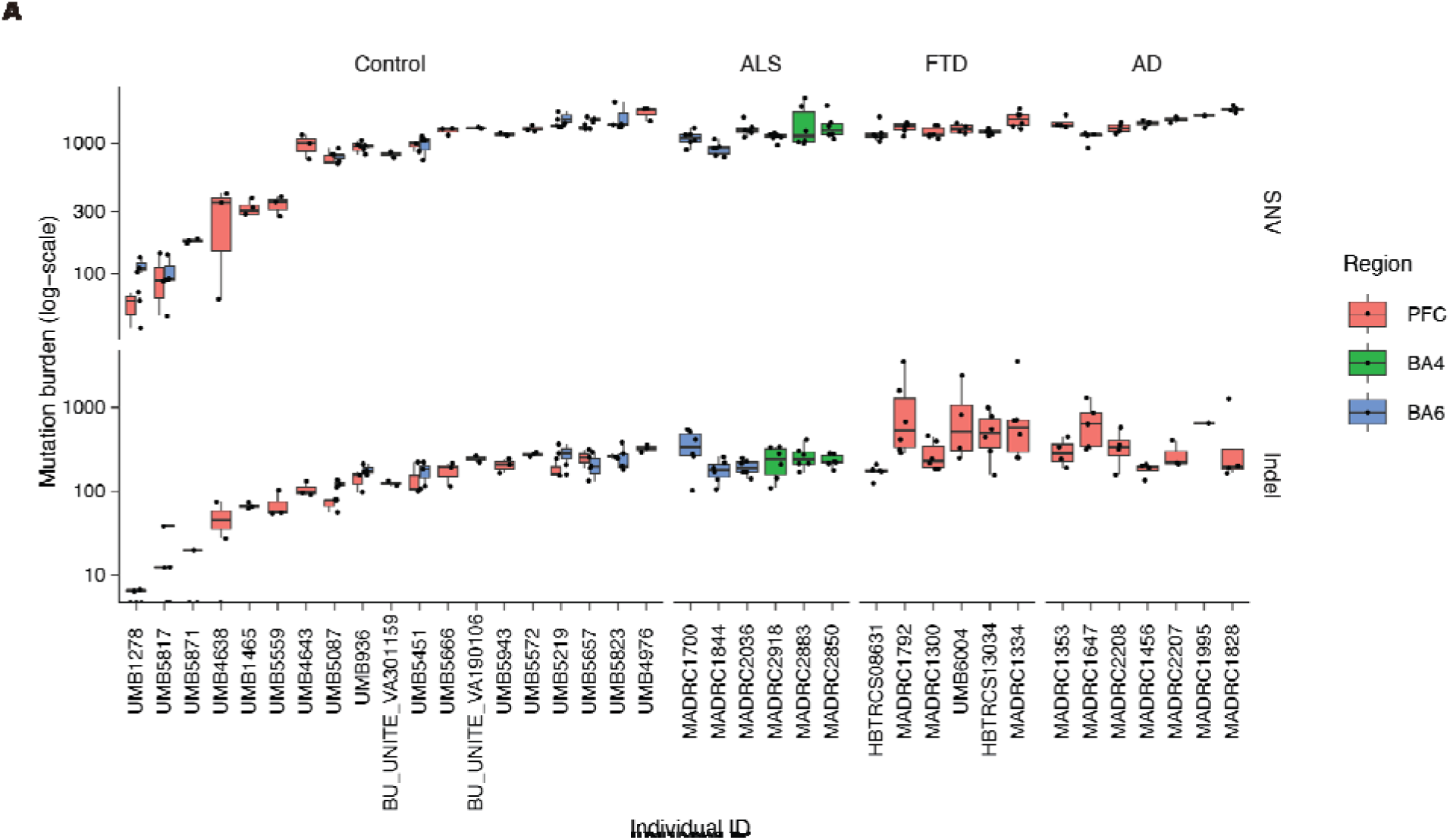
Somatic Indel burdens show large intra-individual variance, related to Figure 2. (A) Burdens of sSNVs and sIndels in PTA-amplified neurons grouped by individual case. Colors indicate brain regions (red: prefrontal cortex; green: primary motor cortex; blue: premotor cortex). Mutation burdens are plotted on a logarithmic scale.

**Data S2.**
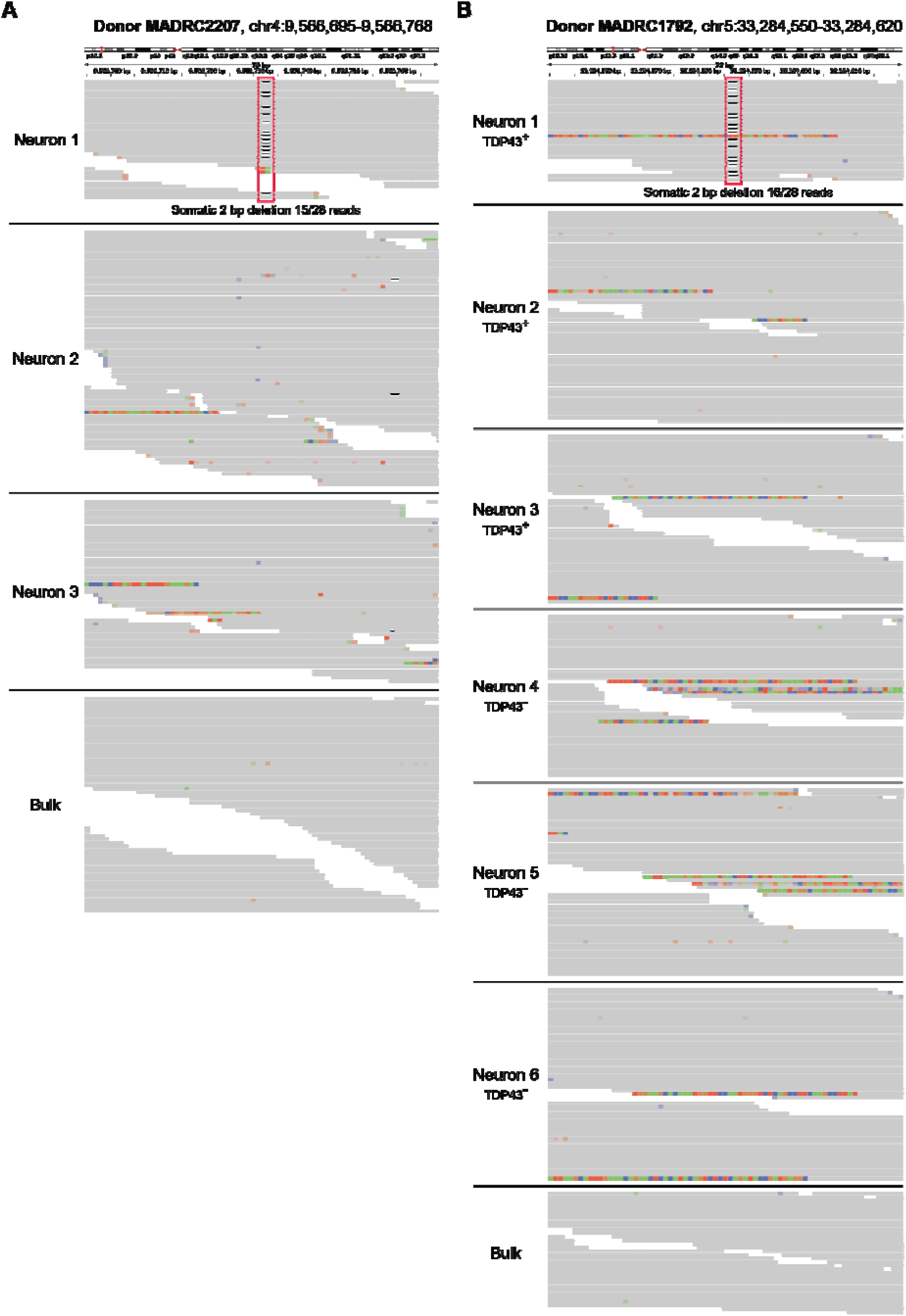
Representative IGV plots of example 2 bp deletions, related to Figure 3. (A) Read pileups from three PTA-amplified neurons and the matched bulk from an individual with AD. (B) Read pileups from six PTA-amplified neurons (TDP-43+ and TDP-43-) and the matched bulk from an individual with FTD. Deletions were supported only by reads in the sample containing the mutation call.

## References

1. Niccoli, T., Partridge, L., and Isaacs, A.M. (2017). Ageing as a risk factor for ALS/FTD. Hum Mol Genet 26, R105–R113. 10.1093/hmg/ddx247.

2. Hebert, L.E., Scherr, P.A., Beckett, L.A., Albert, M.S., Pilgrim, D.M., Chown, M.J., Funkenstein, H.H., and Evans, D.A. (1995). Age-specific incidence of Alzheimer’s disease in a community population. JAMA 273, 1354–1359.

3. Ling, S.C., Polymenidou, M., and Cleveland, D.W. (2013). Converging mechanisms in ALS and FTD: disrupted RNA and protein homeostasis. Neuron 79, 416–438. 10.1016/j.neuron.2013.07.033.

4. DeJesus-Hernandez, M., Mackenzie, I.R., Boeve, B.F., Boxer, A.L., Baker, M., Rutherford, N.J., Nicholson, A.M., Finch, N.A., Flynn, H., Adamson, J., et al. (2011). Expanded GGGGCC hexanucleotide repeat in noncoding region of C9ORF72 causes chromosome 9p-linked FTD and ALS. Neuron 72, 245–256. 10.1016/j.neuron.2011.09.011.

5. Renton, A.E., Majounie, E., Waite, A., Simon-Sanchez, J., Rollinson, S., Gibbs, J.R., Schymick, J.C., Laaksovirta, H., van Swieten, J.C., Myllykangas, L., et al. (2011). A hexanucleotide repeat expansion in C9ORF72 is the cause of chromosome 9p21-linked ALS-FTD. Neuron 72, 257–268. 10.1016/j.neuron.2011.09.010.

6. Hyman, B.T., Phelps, C.H., Beach, T.G., Bigio, E.H., Cairns, N.J., Carrillo, M.C., Dickson, D.W., Duyckaerts, C., Frosch, M.P., Masliah, E., et al. (2012). National Institute on Aging-Alzheimer’s Association guidelines for the neuropathologic assessment of Alzheimer’s disease. Alzheimers Dement 8, 1–13. 10.1016/j.jalz.2011.10.007.

7. Ferrante, R.J., Browne, S.E., Shinobu, L.A., Bowling, A.C., Baik, M.J., MacGarvey, U., Kowall, N.W., Brown, R.H., Jr., and Beal, M.F. (1997). Evidence of increased oxidative damage in both sporadic and familial amyotrophic lateral sclerosis. J Neurochem 69, 2064–2074. 10.1046/j.1471-4159.1997.69052064.x.

8. Haeusler, A.R., Donnelly, C.J., Periz, G., Simko, E.A., Shaw, P.G., Kim, M.S., Maragakis, N.J., Troncoso, J.C., Pandey, A., Sattler, R., et al. (2014). C9orf72 nucleotide repeat structures initiate molecular cascades of disease. Nature 507, 195–200. 10.1038/nature13124.

9. Mitra, J., Guerrero, E.N., Hegde, P.M., Liachko, N.F., Wang, H., Vasquez, V., Gao, J., Pandey, A., Taylor, J.P., Kraemer, B.C., et al. (2019). Motor neuron disease-associated loss of nuclear TDP-43 is linked to DNA double-strand break repair defects. Proc Natl Acad Sci U S A 116, 4696–4705. 10.1073/pnas.1818415116.

10. Smith, C.D., Carney, J.M., Starke-Reed, P.E., Oliver, C.N., Stadtman, E.R., Floyd, R.A., and Markesbery, W.R. (1991). Excess brain protein oxidation and enzyme dysfunction in normal aging and in Alzheimer disease. Proc Natl Acad Sci U S A 88, 10540–10543. 10.1073/pnas.88.23.10540.

11. Adamec, E., Vonsattel, J.P., and Nixon, R.A. (1999). DNA strand breaks in Alzheimer’s disease. Brain Res 849, 67–77. 10.1016/s0006-8993(99)02004-1.

12. Gabbita, S.P., Lovell, M.A., and Markesbery, W.R. (1998). Increased nuclear DNA oxidation in the brain in Alzheimer’s disease. J Neurochem 71, 2034–2040. 10.1046/j.1471-4159.1998.71052034.x.

13. Lodato, M.A., Rodin, R.E., Bohrson, C.L., Coulter, M.E., Barton, A.R., Kwon, M., Sherman, M.A., Vitzthum, C.M., Luquette, L.J., Yandava, C.N., et al. (2018). Aging and neurodegeneration are associated with increased mutations in single human neurons. Science 359, 555–559. 10.1126/science.aao4426.

14. Miller, M.B., Huang, A.Y., Kim, J., Zhou, Z., Kirkham, S.L., Maury, E.A., Ziegenfuss, J.S., Reed, H.C., Neil, J.E., Rento, L., et al. (2022). Somatic genomic changes in single Alzheimer’s disease neurons. Nature 604, 714–722. 10.1038/s41586-022-04640-1.

15. Wang, J., Markesbery, W.R., and Lovell, M.A. (2006). Increased oxidative damage in nuclear and mitochondrial DNA in mild cognitive impairment. J Neurochem 96, 825–832. 10.1111/j.1471-4159.2005.03615.x.

16. Lopez-Gonzalez, R., Lu, Y., Gendron, T.F., Karydas, A., Tran, H., Yang, D., Petrucelli, L., Miller, B.L., Almeida, S., and Gao, F.B. (2016). Poly(GR) in C9ORF72-Related ALS/FTD Compromises Mitochondrial Function and Increases Oxidative Stress and DNA Damage in iPSC-Derived Motor Neurons. Neuron 92, 383–391. 10.1016/j.neuron.2016.09.015.

17. Rosen, D.R., Siddique, T., Patterson, D., Figlewicz, D.A., Sapp, P., Hentati, A., Donaldson, D., Goto, J., O’Regan, J.P., Deng, H.X., and, et al. (1993). Mutations in Cu/Zn superoxide dismutase gene are associated with familial amyotrophic lateral sclerosis. Nature 362, 59–62. 10.1038/362059a0.

18. Acs, K., Luijsterburg, M.S., Ackermann, L., Salomons, F.A., Hoppe, T., and Dantuma, N.P. (2011). The AAA-ATPase VCP/p97 promotes 53BP1 recruitment by removing L3MBTL1 from DNA double-strand breaks. Nat Struct Mol Biol 18, 1345–1350. 10.1038/nsmb.2188.

19. Meerang, M., Ritz, D., Paliwal, S., Garajova, Z., Bosshard, M., Mailand, N., Janscak, P., Hubscher, U., Meyer, H., and Ramadan, K. (2011). The ubiquitin-selective segregase VCP/p97 orchestrates the response to DNA double-strand breaks. Nat Cell Biol 13, 1376–1382. 10.1038/ncb2367.

20. Wang, W.Y., Pan, L., Su, S.C., Quinn, E.J., Sasaki, M., Jimenez, J.C., Mackenzie, I.R., Huang, E.J., and Tsai, L.H. (2013). Interaction of FUS and HDAC1 regulates DNA damage response and repair in neurons. Nat Neurosci 16, 1383–1391. 10.1038/nn.3514.

21. Hewitt, G., Carroll, B., Sarallah, R., Correia-Melo, C., Ogrodnik, M., Nelson, G., Otten, E.G., Manni, D., Antrobus, R., Morgan, B.A., et al. (2016). SQSTM1/p62 mediates crosstalk between autophagy and the UPS in DNA repair. Autophagy 12, 1917–1930. 10.1080/15548627.2016.1210368.

22. Higelin, J., Catanese, A., Semelink-Sedlacek, L.L., Oeztuerk, S., Lutz, A.K., Bausinger, J., Barbi, G., Speit, G., Andersen, P.M., Ludolph, A.C., et al. (2018). NEK1 loss-of-function mutation induces DNA damage accumulation in ALS patient-derived motoneurons. Stem Cell Res 30, 150–162. 10.1016/j.scr.2018.06.005.

23. Cohen, S., Puget, N., Lin, Y.L., Clouaire, T., Aguirrebengoa, M., Rocher, V., Pasero, P., Canitrot, Y., and Legube, G. (2018). Senataxin resolves RNA:DNA hybrids forming at DNA double-strand breaks to prevent translocations. Nat Commun 9, 533. 10.1038/s41467-018-02894-w.

24. Maor-Nof, M., Shipony, Z., Lopez-Gonzalez, R., Nakayama, L., Zhang, Y.J., Couthouis, J., Blum, J.A., Castruita, P.A., Linares, G.R., Ruan, K., et al. (2021). p53 is a central regulator driving neurodegeneration caused by C9orf72 poly(PR). Cell 184, 689–708 e620. 10.1016/j.cell.2020.12.025.

25. Mirceta, M., Schmidt, M.H.M., Shum, N., Prasolava, T.K., Meikle, B., Lanni, S., Mohiuddin, M., McKeever, P.M., Zhang, M., Liang, M., et al. (2024). C9orf72 repeat expansion creates the unstable folate-sensitive fragile site FRA9A. NAR Mol Med 1, ugae019. 10.1093/narmme/ugae019.

26. Luquette, L.J., Miller, M.B., Zhou, Z., Bohrson, C.L., Zhao, Y., Jin, H., Gulhan, D., Ganz, J., Bizzotto, S., Kirkham, S., et al. (2022). Single-cell genome sequencing of human neurons identifies somatic point mutation and indel enrichment in regulatory elements. Nat Genet 54, 1564–1571. 10.1038/s41588-022-01180-2.

27. Liu, E.Y., Russ, J., Cali, C.P., Phan, J.M., Amlie-Wolf, A., and Lee, E.B. (2019). Loss of Nuclear TDP-43 Is Associated with Decondensation of LINE Retrotransposons. Cell Rep 27, 1409–1421 e1406. 10.1016/j.celrep.2019.04.003.

28. Ma, X.R., Prudencio, M., Koike, Y., Vatsavayai, S.C., Kim, G., Harbinski, F., Briner, A., Rodriguez, C.M., Guo, C., Akiyama, T., et al. (2022). TDP-43 represses cryptic exon inclusion in the FTD-ALS gene UNC13A. Nature 603, 124–130. 10.1038/s41586-022-04424-7.

29. Gittings, L.M., Alsop, E.B., Antone, J., Singer, M., Whitsett, T.G., Sattler, R., and Van Keuren-Jensen, K. (2023). Cryptic exon detection and transcriptomic changes revealed in single-nuclei RNA sequencing of C9ORF72 patients spanning the ALS-FTD spectrum. Acta Neuropathol 146, 433–450. 10.1007/s00401-023-02599-5.

30. Ling, J.P., Pletnikova, O., Troncoso, J.C., and Wong, P.C. (2015). TDP-43 repression of nonconserved cryptic exons is compromised in ALS-FTD. Science 349, 650–655. 10.1126/science.aab0983.

31. Nolan, M., Scott, C., Gamarallage, M.P., Lunn, D., Carpenter, K., McDonough, E., Meyer, D., Kaanumalle, S., Santamaria-Pang, A., Turner, M.R., et al. (2020). Quantitative patterns of motor cortex proteinopathy across ALS genotypes. Acta Neuropathol Commun 8, 98. 10.1186/s40478-020-00961-2.

32. Foerster, B.R., Pomper, M.G., Callaghan, B.C., Petrou, M., Edden, R.A., Mohamed, M.A., Welsh, R.C., Carlos, R.C., Barker, P.B., and Feldman, E.L. (2013). An imbalance between excitatory and inhibitory neurotransmitters in amyotrophic lateral sclerosis revealed by use of 3-T proton magnetic resonance spectroscopy. JAMA Neurol 70, 1009–1016. 10.1001/jamaneurol.2013.234.

33. Khademullah, C.S., Aqrabawi, A.J., Place, K.M., Dargaei, Z., Liang, X., Pressey, J.C., Bedard, S., Yang, J.W., Garand, D., Keramidis, I., et al. (2020). Cortical interneuron-mediated inhibition delays the onset of amyotrophic lateral sclerosis. Brain 143, 800–810. 10.1093/brain/awaa034.

34. Allodi, I., Montanana-Rosell, R., Selvan, R., Low, P., and Kiehn, O. (2021). Locomotor deficits in a mouse model of ALS are paralleled by loss of V1-interneuron connections onto fast motor neurons. Nat Commun 12, 3251. 10.1038/s41467-021-23224-7.

35. Ganz, J., Luquette, L.J., Bizzotto, S., Miller, M.B., Zhou, Z., Bohrson, C.L., Jin, H., Tran, A.V., Viswanadham, V.V., McDonough, G., et al. (2024). Contrasting somatic mutation patterns in aging human neurons and oligodendrocytes. Cell 187, 1955–1970 e1923. 10.1016/j.cell.2024.02.025.

36. Affymetrix (2008). Affymetrix® White Paper: Median of the Absolute Values of all Pairwise Differences and Quality Control on the Affymetrix Genome-Wide Human SNP Array 6.0

37. Charcot, J.M. (1880). De la sclerose laterale amyotrophique. Symptomatologie. Lecons sur les Maladies du Systeme Nerveux Faites a la Salpetriere 2, 227–242.

38. Alexandrov, L.B., Nik-Zainal, S., Wedge, D.C., Campbell, P.J., and Stratton, M.R. (2013). Deciphering signatures of mutational processes operative in human cancer. Cell Rep 3, 246–259. 10.1016/j.celrep.2012.12.008.

39. Petljak, M., Alexandrov, L.B., Brammeld, J.S., Price, S., Wedge, D.C., Grossmann, S., Dawson, K.J., Ju, Y.S., Iorio, F., Tubio, J.M.C., et al. (2019). Characterizing Mutational Signatures in Human Cancer Cell Lines Reveals Episodic APOBEC Mutagenesis. Cell 176, 1282–1294 e1220. 10.1016/j.cell.2019.02.012.

40. Klungland, A., Hoss, M., Gunz, D., Constantinou, A., Clarkson, S.G., Doetsch, P.W., Bolton, P.H., Wood, R.D., and Lindahl, T. (1999). Base excision repair of oxidative DNA damage activated by XPG protein. Mol Cell 3, 33–42. 10.1016/s1097-2765(00)80172-0.

41. Reijns, M.A.M., Parry, D.A., Williams, T.C., Nadeu, F., Hindshaw, R.L., Rios Szwed, D.O., Nicholson, M.D., Carroll, P., Boyle, S., Royo, R., et al. (2022). Signatures of TOP1 transcription-associated mutagenesis in cancer and germline. Nature 602, 623–631. 10.1038/s41586-022-04403-y.

42. Alexandrov, L.B., Kim, J., Haradhvala, N.J., Huang, M.N., Tian Ng, A.W., Wu, Y., Boot, A., Covington, K.R., Gordenin, D.A., Bergstrom, E.N., et al. (2020). The repertoire of mutational signatures in human cancer. Nature 578, 94–101. 10.1038/s41586-020-1943-3.

43. Tanizawa, A., Kohn, K.W., and Pommier, Y. (1993). Induction of cleavage in topoisomerase I c-DNA by topoisomerase I enzymes from calf thymus and wheat germ in the presence and absence of camptothecin. Nucleic Acids Res 21, 5157–5166. 10.1093/nar/21.22.5157.

44. Kiianitsa, K., and Maizels, N. (2013). A rapid and sensitive assay for DNA-protein covalent complexes in living cells. Nucleic Acids Res 41, e104. 10.1093/nar/gkt171.

45. Pommier, Y., Sun, Y., Huang, S.N., and Nitiss, J.L. (2016). Roles of eukaryotic topoisomerases in transcription, replication and genomic stability. Nat Rev Mol Cell Biol 17, 703–721. 10.1038/nrm.2016.111.

46. Roadmap Epigenomics, C., Kundaje, A., Meuleman, W., Ernst, J., Bilenky, M., Yen, A., Heravi-Moussavi, A., Kheradpour, P., Zhang, Z., Wang, J., et al. (2015). Integrative analysis of 111 reference human epigenomes. Nature 518, 317–330. 10.1038/nature14248.

47. Ernst, J., and Kellis, M. (2017). Chromatin-state discovery and genome annotation with ChromHMM. Nat Protoc 12, 2478–2492. 10.1038/nprot.2017.124.

48. Nott, A., Holtman, I.R., Coufal, N.G., Schlachetzki, J.C.M., Yu, M., Hu, R., Han, C.Z., Pena, M., Xiao, J., Wu, Y., et al. (2019). Brain cell type-specific enhancer-promoter interactome maps and disease-risk association. Science 366, 1134–1139. 10.1126/science.aay0793.

49. Xing, D., Tan, L., Chang, C.H., Li, H., and Xie, X.S. (2021). Accurate SNV detection in single cells by transposon-based whole-genome amplification of complementary strands. Proc Natl Acad Sci U S A 118. 10.1073/pnas.2013106118.

50. Gronska-Peski, M., Srinivasa, A., and Evrony, G.D. (2025). Divergent somatic mutation patterns among human cerebellar neuron types. bioRxiv. 10.1101/2025.09.29.679392.

51. Essuman, K., Yang, Y., Goodman, E., Cambridge, C.N., Cai, C., An, Z., Mao, S., Manam, M.D., Finander, B., Khoshkhoo, S., et al. (2026). Somatic mutation in human cerebellum illustrates neuron type-specific patterns of age-related mutation. bioRxiv. 10.64898/2026.02.27.708647.

52. Cerritelli, S.M., and Crouch, R.J. (2016). The Balancing Act of Ribonucleotides in DNA. Trends Biochem Sci 41, 434–445. 10.1016/j.tibs.2016.02.005.

53. Crespan, E., Furrer, A., Rosinger, M., Bertoletti, F., Mentegari, E., Chiapparini, G., Imhof, R., Ziegler, N., Sturla, S.J., Hubscher, U., et al. (2016). Impact of ribonucleotide incorporation by DNA polymerases beta and lambda on oxidative base excision repair. Nat Commun 7, 10805. 10.1038/ncomms10805.

54. Pourquier, P., Ueng, L.M., Fertala, J., Wang, D., Park, H.J., Essigmann, J.M., Bjornsti, M.A., and Pommier, Y. (1999). Induction of reversible complexes between eukaryotic DNA topoisomerase I and DNA-containing oxidative base damages. 7, 8-dihydro-8-oxoguanine and 5-hydroxycytosine. J Biol Chem 274, 8516–8523. 10.1074/jbc.274.13.8516.

55. Pommier, Y., Nussenzweig, A., Takeda, S., and Austin, C. (2022). Human topoisomerases and their roles in genome stability and organization. Nat Rev Mol Cell Biol 23, 407–427. 10.1038/s41580-022-00452-3.

56. Crewe, M., and Madabhushi, R. (2021). Topoisomerase-Mediated DNA Damage in Neurological Disorders. Front Aging Neurosci 13, 751742. 10.3389/fnagi.2021.751742.

57. Jeffries, A.M., Yu, T., Ziegenfuss, J.S., Tolles, A.K., Baer, C.E., Sotelo, C.B., Kim, Y., Weng, Z., and Lodato, M.A. (2025). Single-cell transcriptomic and genomic changes in the ageing human brain. Nature 646, 657–666. 10.1038/s41586-025-09435-8.

58. Caldecott, K.W., Ward, M.E., and Nussenzweig, A. (2022). The threat of programmed DNA damage to neuronal genome integrity and plasticity. Nat Genet 54, 115–120. 10.1038/s41588-021-01001-y.

59. King, I.F., Yandava, C.N., Mabb, A.M., Hsiao, J.S., Huang, H.S., Pearson, B.L., Calabrese, J.M., Starmer, J., Parker, J.S., Magnuson, T., et al. (2013). Topoisomerases facilitate transcription of long genes linked to autism. Nature 501, 58–62. 10.1038/nature12504.

60. Zylka, M.J., Simon, J.M., and Philpot, B.D. (2015). Gene length matters in neurons. Neuron 86, 353–355. 10.1016/j.neuron.2015.03.059.

61. Fragola, G., Mabb, A.M., Taylor-Blake, B., Niehaus, J.K., Chronister, W.D., Mao, H., Simon, J.M., Yuan, H., Li, Z., McConnell, M.J., and Zylka, M.J. (2020). Deletion of Topoisomerase 1 in excitatory neurons causes genomic instability and early onset neurodegeneration. Nat Commun 11, 1962. 10.1038/s41467-020-15794-9.

62. Fielden, J., Wiseman, K., Torrecilla, I., Li, S., Hume, S., Chiang, S.C., Ruggiano, A., Narayan Singh, A., Freire, R., Hassanieh, S., et al. (2020). TEX264 coordinates p97- and SPRTN-mediated resolution of topoisomerase 1-DNA adducts. Nat Commun 11, 1274. 10.1038/s41467-020-15000-w.

63. Lascaux, P., Hoslett, G., Tribble, S., Trugenberger, C., Anticevic, I., Otten, C., Torrecilla, I., Koukouravas, S., Zhao, Y., Yang, H., et al. (2024). TEX264 drives selective autophagy of DNA lesions to promote DNA repair and cell survival. Cell 187, 5698–5718 e5626. 10.1016/j.cell.2024.08.020.

64. Dong, G., Ma, C.C., Mao, S., Naik, S.M., Brown, K.S.-M., McDonough, G.A., Kim, J., Kirkham, S.L., Cherry, J.D., Uretsky, M., et al. (2025). Diverse somatic genomic alterations in single neurons in chronic traumatic encephalopathy. bioRxiv, 2025.2003.2003.641217. 10.1101/2025.03.03.641217.

65. McKee, A.C., Stein, T.D., Kiernan, P.T., and Alvarez, V.E. (2015). The neuropathology of chronic traumatic encephalopathy. Brain Pathol 25, 350–364. 10.1111/bpa.12248.

66. Josephs, K.A., Murray, M.E., Whitwell, J.L., Parisi, J.E., Petrucelli, L., Jack, C.R., Petersen, R.C., and Dickson, D.W. (2014). Staging TDP-43 pathology in Alzheimer’s disease. Acta Neuropathol 127, 441–450. 10.1007/s00401-013-1211-9.

67. McKee, A.C., Gavett, B.E., Stern, R.A., Nowinski, C.J., Cantu, R.C., Kowall, N.W., Perl, D.P., Hedley-Whyte, E.T., Price, B., Sullivan, C., et al. (2010). TDP-43 proteinopathy and motor neuron disease in chronic traumatic encephalopathy. J Neuropathol Exp Neurol 69, 918–929. 10.1097/NEN.0b013e3181ee7d85.

68. Kulbe, J.R., and Hall, E.D. (2017). Chronic traumatic encephalopathy-integration of canonical traumatic brain injury secondary injury mechanisms with tau pathology. Prog Neurobiol 158, 15–44. 10.1016/j.pneurobio.2017.08.003.

69. Sugawara, S., Okada, R., Loo, T.M., Tanaka, H., Miyata, K., Chiba, M., Kawasaki, H., Katoh, K., Kaji, S., Maezawa, Y., et al. (2022). RNaseH2A downregulation drives inflammatory gene expression via genomic DNA fragmentation in senescent and cancer cells. Commun Biol 5, 1420. 10.1038/s42003-022-04369-7.

70. Gao, C., Jiang, J., Tan, Y., and Chen, S. (2023). Microglia in neurodegenerative diseases: mechanism and potential therapeutic targets. Signal Transduct Target Ther 8, 359. 10.1038/s41392-023-01588-0.

71. Crow, Y.J., Leitch, A., Hayward, B.E., Garner, A., Parmar, R., Griffith, E., Ali, M., Semple, C., Aicardi, J., Babul-Hirji, R., et al. (2006). Mutations in genes encoding ribonuclease H2 subunits cause Aicardi-Goutieres syndrome and mimic congenital viral brain infection. Nat Genet 38, 910–916. 10.1038/ng1842.

72. Katyal, S., Lee, Y., Nitiss, K.C., Downing, S.M., Li, Y., Shimada, M., Zhao, J., Russell, H.R., Petrini, J.H., Nitiss, J.L., and McKinnon, P.J. (2014). Aberrant topoisomerase-1 DNA lesions are pathogenic in neurodegenerative genome instability syndromes. Nat Neurosci 17, 813–821. 10.1038/nn.3715.

73. Zhao, X.N., and Usdin, K. (2015). The Repeat Expansion Diseases: The dark side of DNA repair. DNA Repair (Amst) 32, 96–105. 10.1016/j.dnarep.2015.04.019.

74. Walton, R.T., Qin, Y., Kirby, B., Andrews, J.O., Taipale, M., Kang, B., and Blainey, P.C. (2025). CROPseq-multi: a universal solution for multiplexed perturbation in high-content pooled CRISPR screens. bioRxiv. 10.1101/2024.03.17.585235.

75. Evrony, G.D., Cai, X., Lee, E., Hills, L.B., Elhosary, P.C., Lehmann, H.S., Parker, J.J., Atabay, K.D., Gilmore, E.C., Poduri, A., et al. (2012). Single-neuron sequencing analysis of L1 retrotransposition and somatic mutation in the human brain. Cell 151, 483–496. 10.1016/j.cell.2012.09.035.

76. Bergstrom, E.N., Huang, M.N., Mahto, U., Barnes, M., Stratton, M.R., Rozen, S.G., and Alexandrov, L.B. (2019). SigProfilerMatrixGenerator: a tool for visualizing and exploring patterns of small mutational events. BMC Genomics 20, 685. 10.1186/s12864-019-6041-2.

77. Islam, S.M.A., Diaz-Gay, M., Wu, Y., Barnes, M., Vangara, R., Bergstrom, E.N., He, Y., Vella, M., Wang, J., Teague, J.W., et al. (2022). Uncovering novel mutational signatures by de novo extraction with SigProfilerExtractor. Cell Genom 2, None. 10.1016/j.xgen.2022.100179.

78. Consortium, G.T. (2020). The GTEx Consortium atlas of genetic regulatory effects across human tissues. Science 369, 1318–1330. 10.1126/science.aaz1776.

79. Cotto, K.C., Feng, Y.Y., Ramu, A., Richters, M., Freshour, S.L., Skidmore, Z.L., Xia, H., McMichael, J.F., Kunisaki, J., Campbell, K.M., et al. (2023). Integrated analysis of genomic and transcriptomic data for the discovery of splice-associated variants in cancer. Nat Commun 14, 1589. 10.1038/s41467-023-37266-6.

80. Li, Y.I., Knowles, D.A., Humphrey, J., Barbeira, A.N., Dickinson, S.P., Im, H.K., and Pritchard, J.K. (2018). Annotation-free quantification of RNA splicing using LeafCutter. Nat Genet 50, 151–158. 10.1038/s41588-017-0004-9.

81. Li, H. (2013). Aligning sequence reads, clone sequences and assembly contigs with BWA-MEM. arXiv preprint arXiv:1303.3997.

82. Dong, G., Hilal, N., Mallett, S., Jin, B., Mao, S., Manam, M.D., Shao, D.D., Choudhury, S., Huang, A.Y., and Lee, E.A. (2026). Duplex-Indel: a snakemake pipeline for somatic indel calling in Tn5 transposase-based duplex sequencing data. Bioinformatics. 10.1093/bioinformatics/btag205.

83. Zheng, G.X., Terry, J.M., Belgrader, P., Ryvkin, P., Bent, Z.W., Wilson, R., Ziraldo, S.B., Wheeler, T.D., McDermott, G.P., Zhu, J., et al. (2017). Massively parallel digital transcriptional profiling of single cells. Nat Commun 8, 14049. 10.1038/ncomms14049.

84. Fleming, S.J., Chaffin, M.D., Arduini, A., Akkad, A.D., Banks, E., Marioni, J.C., Philippakis, A.A., Ellinor, P.T., and Babadi, M. (2023). Unsupervised removal of systematic background noise from droplet-based single-cell experiments using CellBender. Nat Methods 20, 1323–1335. 10.1038/s41592-023-01943-7.

85. Germain, P.L., Lun, A., Garcia Meixide, C., Macnair, W., and Robinson, M.D. (2021). Doublet identification in single-cell sequencing data using scDblFinder. F1000Res 10, 979. 10.12688/f1000research.73600.2.

86. Muskovic, W., and Powell, J.E. (2021). DropletQC: improved identification of empty droplets and damaged cells in single-cell RNA-seq data. Genome Biol 22, 329. 10.1186/s13059-021-02547-0.

87. Hao, Y., Stuart, T., Kowalski, M.H., Choudhary, S., Hoffman, P., Hartman, A., Srivastava, A., Molla, G., Madad, S., Fernandez-Granda, C., and Satija, R. (2024). Dictionary learning for integrative, multimodal and scalable single-cell analysis. Nat Biotechnol 42, 293–304. 10.1038/s41587-023-01767-y.

88. Korsunsky, I., Millard, N., Fan, J., Slowikowski, K., Zhang, F., Wei, K., Baglaenko, Y., Brenner, M., Loh, P.R., and Raychaudhuri, S. (2019). Fast, sensitive and accurate integration of single-cell data with Harmony. Nat Methods 16, 1289–1296. 10.1038/s41592-019-0619-0.

89. Traag, V.A., Waltman, L., and van Eck, N.J. (2019). From Louvain to Leiden: guaranteeing well-connected communities. Sci Rep 9, 5233. 10.1038/s41598-019-41695-z.

90. Bakken, T.E., Jorstad, N.L., Hu, Q., Lake, B.B., Tian, W., Kalmbach, B.E., Crow, M., Hodge, R.D., Krienen, F.M., Sorensen, S.A., et al. (2021). Comparative cellular analysis of motor cortex in human, marmoset and mouse. Nature 598, 111–119. 10.1038/s41586-021-03465-8.

91. Lai, J., Demirbas, D., Kim, J., Jeffries, A.M., Tolles, A., Park, J., Chittenden, T.W., Buckley, P.G., Yu, T.W., Lodato, M.A., and Lee, E.A. (2024). ATM-deficiency-induced microglial activation promotes neurodegeneration in ataxia-telangiectasia. Cell Rep 43, 113622. 10.1016/j.celrep.2023.113622.

92. Harrison, J.G., Calder, W.J., Shastry, V., and Buerkle, C.A. (2020). Dirichlet-multinomial modelling outperforms alternatives for analysis of microbiome and other ecological count data. Mol Ecol Resour 20, 481–497. 10.1111/1755-0998.13128.

93. He, L., Davila-Velderrain, J., Sumida, T.S., Hafler, D.A., Kellis, M., and Kulminski, A.M. (2021). NEBULA is a fast negative binomial mixed model for differential or co-expression analysis of large-scale multi-subject single-cell data. Commun Biol 4, 629. 10.1038/s42003-021-02146-6.

94. Gabitto, M.I., Travaglini, K.J., Rachleff, V.M., Kaplan, E.S., Long, B., Ariza, J., Ding, Y., Mahoney, J.T., Dee, N., Goldy, J., et al. (2024). Integrated multimodal cell atlas of Alzheimer’s disease. Nat Neurosci 27, 2366–2383. 10.1038/s41593-024-01774-5.

95. Fernandopulle, M.S., Prestil, R., Grunseich, C., Wang, C., Gan, L., and Ward, M.E. (2018). Transcription Factor-Mediated Differentiation of Human iPSCs into Neurons. Curr Protoc Cell Biol 79, e51. 10.1002/cpcb.51.

